# Th17 cells target the metabolic miR-142-5p-SDHC/SDHD axis promoting invasiveness and progression of cervical cancers

**DOI:** 10.1101/2023.06.29.547020

**Authors:** Maike Pohlers, Selina Gies, Tanja Tänzer, Russalina Stroeder, Laura Theobald, Nicole Ludwig, Yoo-Jin Kim, Rainer Bohle, Erich-Franz Solomayer, Eckart Meese, Martin Hart, Barbara Walch-Rückheim

**Affiliations:** Center of Human and Molecular Biology (ZHMB), Institute of Virology, Saarland University, Homburg/Saar, Germany; Department of Obstetrics and Gynecology, Saarland University Medical Center, Homburg/Saar, Germany; Institute of Human Genetics, Saarland University, Homburg/Saar, Germany; Institute of Pathology, Saarland University Medical Center, Homburg/Saar, Germany

**Keywords:** T-helper-17 cells, cervical cancers, miR-142-5p, succinate dehydrogenase complex, metastases

## Abstract

During cervical carcinogenesis, T-helper (Th)-17 cells accumulate in the peripheral blood and tumor tissues of cancer patients. We previously demonstrated that Th17 cells are associated with therapy resistance as well as cervical cancer metastases and relapse, however, the underlying Th17-driven mechanisms supporting cervical cancer progression are not fully understood as yet. In this study, we found that Th17 cells promote migration and invasion of cervical cancer cells in 2D cultures and 3D spheroids. We demonstrated that Th17 cells induced the expression of miR-142-5p in cervical cancer cells supporting their migration and invasiveness. As the responsible mechanism, we identified the subunits C and D of the succinate dehydrogenase (SDH) complex as new targets of miR-142-5p and provided evidence that Th17 cells reduced the expression of SDHC and SDHD that was dependent on miR-142-5p. Functional downstream analysis with inhibitors of miR-142-5p and siRNA knock down of SDHC and SDHD revealed that Th17-induced miR-142-5p-mediated reduced expression of SDHC and SDHD was responsible for enhanced migration and invasion of cervical cancer cells. Consistently, cervical cancer patients exhibited high levels of succinate in their serum associated with lymph node metastases and diminished expression of SDHD in patients′ biopsies significantly correlated with increased numbers of Th17 cells, advanced tumor stage and lymph node metastases. Correspondingly, a combination of weak or negative SDHD expression and a ratio of Th17/CD4^+^ T cells > 43.90 % *in situ* was associated with reduced recurrence free survival. In summary, we unraveled a novel molecular mechanism by which Th17 cells promote cervical cancer progression and suggest evaluation of Th17 cells as a potential target for immunotherapy in cervical cancer.

## Introduction

Cervical cancer is the fourth most common cancer in woman worldwide and the consequence of persistent infection of high risk human papillomaviruses (HPV) (1). Treatment approaches depend on the disease stages according to the International Federation of Gynecology and Obstetrics (FIGO) classification, and include besides surgery alone in early stages, adjuvant platinum-based concurrent chemoradiotherapy (CRT) in addition to surgery for advanced cervical cancers with prognostic risk factors for local relapse and primary CRT for locally advanced often inoperable cervical carcinoma (FIGO > IIB) (2). However, depending on initial tumor stage, 8 to 26 % of women with cervical cancer experience relapse of disease, most commonly within the first 2 years of completing primary treatment (3) resulting in 5-year survival raging from over 90 % if diagnosed in an early, localized stage to less than 20 % if diagnosed as distant or metastatic (4). Thus, metastatic cervical cancer remains a major clinical challenge.

The constitution of the HPV-associated tumor microenvironment is critical for tumor development, therapy responses and affects progression of disease (5, 6). Invasive cervical cancers are often associated with strong inflammatory infiltrates in the stroma (7). HPV-transformed keratinocytes actively contribute to the inflammatory microenvironment during cervical carcinogenesis via production of the cytokine interleukin (IL)-6 (8–10) and favor the infiltration of T-helper (Th)17 cells (11). Th17 cells represent an IL-17-expressing subgroup of Th cells (12). They increase in the blood of cervical cancer patients (13), infiltrate cervical cancer tissues (11) and exhibit pro-inflammatory as well as tumor-promoting properties in different cancer types (6, 14, 15). During cervical carcinogenesis, we previously demonstrated that cervical cancer cells actively contributed to the infiltration (11) and expansion (10) of Th17 cells in tumor tissues and their presence *in situ* correlated with advanced tumor stage, lymph node metastases (10, 11), reduced responsiveness of patients toward CRT (6) and retrospectively, with cervical cancer recurrence (6, 10). However, Th17-induced molecular mechanisms favoring cervical cancer metastases and progression are less studied so far.

Besides the role of the tumor micromilieu for cancer development and therapeutic outcome, metabolic changes within tumor cells also affect therapy responses and cancer progression (16). One reason for aberrant metabolic reactions could be a dysfunction of the succinate dehydrogenase (SDH) complex. This complex has a central role in energy metabolism, as it directly links the TCA cycle to the respiratory machinery as complex II. It is composed of four subunits (A, B, C, D). SDHA and B are hydrophilic subunits and form the catalytic unit of the complex, whereas SDHC and SDHD represent the hydrophobic membrane-bound part of the complex (17). Mutations in SDH subunits are linked to familial paraganglioma syndromes, pheochromocytomas, renal cancers and gastrointestinal stromal tumors (18, 19) and reduced expression of subunits B or C supported epithelial-mesenchymal transition (EMT) in breast cancer cells (19), aerobic glycolysis in colon cancer cells (20) and the Warburg effect in hepatocellular carcinomas (21).

In this study, we investigated how Th17 cells contributed to cervical cancer progression. We demonstrated that Th17 cells increased proliferation and migration as well as invasiveness of cervical cancer cells. Our data provide evidence that Th17 cells induced the expression of miR-142-5p in cervical cancer cells and identified SDHC and SDHD as new targets of miR-142-5p responsible for enhanced migration and invasion. Notably, we found that high numbers of Th17 cells correlated with reduced SDHD expression, showing strongest Th17-mediated reduction in expression, in cervical cancer patients *in situ* that were associated with reduced recurrence free survival and high serum levels of succinate were linked with lymph node metastases.

## Results

### Th17 cells induced an EMT phenotype in cervical cancer cells and promoted their migration and invasiveness

Th17 cells infiltrate cervical cancer tissues and are associated with lymph node metastases and cervical cancer recurrence (10, 11). We were interested how Th17 cells might support cervical cancer progression and performed a mRNA expression analysis (Agilent SurePrint G3 Human GE v3 8×60K microarray) of SiHa and SW756 cells after stimulation with rhIL-17, CM of *in vitro* generated Th17 cells or after direct cell-cell contact in co-cultures with Th17 cells (scheme in Supplementary Fig. S1A) Out of 58000 detectable mRNAs of the microarray, 1802 protein coding transcripts were at least 1.5-fold changed after stimulation with rhIL-17, 5639 protein coding transcripts after CM Th17 stimulation and 5162 after co-cultures in comparison to medium stimulated control cells. Analyzing a signature of genes representing an EMT phenotype we found reduced expression of *CDH1* and *CDH2* after co-cultures (Supplementary Fig. S1B), rhIL-17 and CM Th17 stimulation in SiHa and SW756 cells (Fig. 1A). In contrast, the expression of *TWIST1, SNAI1*, *SNAI2*, *ZEB1*, *ZEB2*, *VIM* and *CTNNB1* was increased. Furthermore, Th17 cells induced the expression of members of the S100 family (*S100A2*, *S100A4*, *S100A6*, *S100A8*, *S100A9* and *S100A14*) in co-cultured cervical cancer cells involved in EMT and metastasis (22). Interestingly, induction by rhIL-17 or Th17-produced soluble factors in the CM was stronger (up to fold change >3). Quantitative real time PCRs of four genes were performed to validate the results showing reduced expression of *CDH1* (blue bars) and increased expression of *VIM*, *TWIST1* and *ZEB1* (light and dark red bars) after rhIL-17 and CM Th17 stimulation (Fig. 1B, upper pannel) and co-cultures (Supplementary Fig. S1C). On protein levels, CM of Th17 cells reduced e-cadherin, but enhanced vimentin expression (Fig. 1B, lower panel).

**Figure 1:**
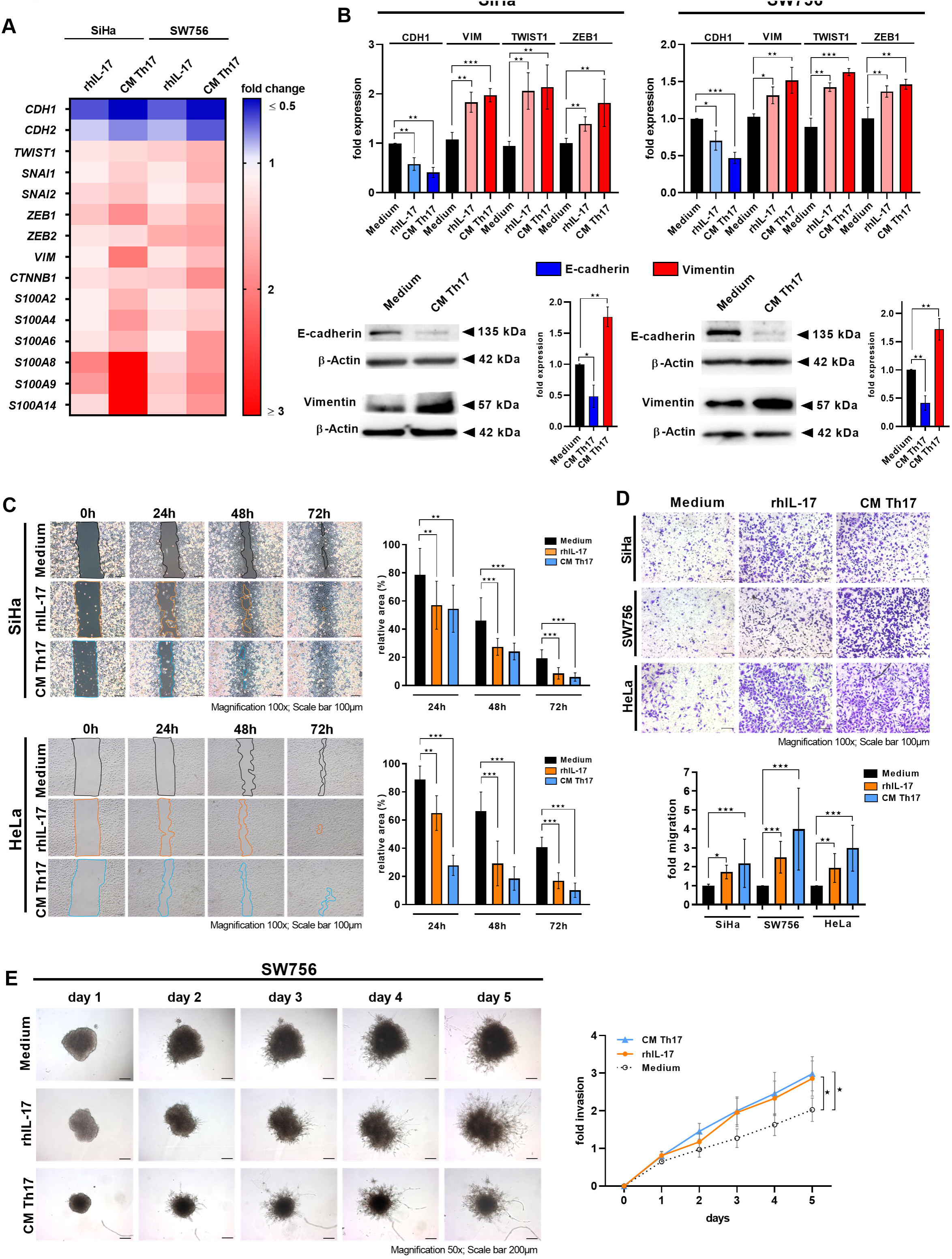
Th17 cells induced the expression of EMT markers in cervical cancer cells and enhanced their migration and invasiveness. (A, B) SiHa and SW756 cells were stimulated with rhIL-17 or CM of Th17 cells for 24 h. (A) Transcriptomic profile of EMT markers. Fold changes of stimulated cells to unstimulated cells were illustrated by color code (reduction in blue, induction in red). (B) Expression profile of EMT markers was validated by qRT-PCR (B upper panel; rhIL17 stimulation: light red and light blue bars, CM Th17 stimulation: dark red and dark blue bars) and western blot analysis after 72 h (lower panel). β-actin was used as a loading control. Bars represent quantification of n=3 independent experiments (mean ±SD; E-cadherin: blue bars; vimentin: red bars). (C) Monolayers of SiHa and HeLa cells stimulated with medium, rhIL-17 or CM of Th17 cells were scratched. Pictures were taken after 0, 24, 48 and 72 h. The area of the gap, indicated by lines (left panel), was determined in relation to time point 0 h (right panel). Scale bar: 100 μm. Bars represents results (mean ±SD) of n=3 experiments. The area of 0 h was set at 100 %, respectively. (D) SiHa, SW756 and HeLa cells were stimulated with medium, rhIL-17 or CM of Th17 cells and used in transwell migration assays. Transmigrated cells were calculated after 24 h. Representative pictures (upper panel; scale bar: 100 µm); quantification of n=3 experiments with five independent pictures, respectively (mean ±SD) lower panel. The number of medium stimulated cells was set at 1. (E) Spheroids of SW756 cells were generated in the absence or presence of rhIL-17 or CM of Th17 cells. Spheroids were embedded into matrigel, pictures were taken for 5 days and spheroid invasion was calculated. Shown are the results mean ± SD from four independent experiments performed in doubles. P-value according to the nonparametric Kruskal-Wallis test (B, C, D) or Mann-Whitney U-test (E). Asterisks represent statistical significances: *P < 0.05; **P < 0.01; ***P < 0.001.

Our results prompted us to analyze whether Th17 cells affect cervical cancer cell migration and invasion. In scratch assays, measuring both migration and proliferation, the introduced gap was closed significantly faster resulting in a significant decrease of relative area over 72h by two different cervical cancer cells after stimulation with rhIL-17 (orange bars) or CM of Th17 cells (blue bars) in comparison to unstimulated cells (black bars) (Fig. 1C). In transwell migration assays, stimulation of three different cervical cancer cells with rhIL-17 (orange bars) or CM of Th17 cells (blue bars) resulted in 1.7-3.9 fold increased migration in comparison to unstimulated cells (Fig. 1D). Finally, we examined the invasive behavior of cervical cancer cells in spheroid invasion assays using basement-membrane like matrix (Matrigel™). After 5 days, medium stimulated spheroids exhibited marginal branch-like invadopodia emerging into the surrounding matrix (Fig. 1D). In contrast, spheroids stimulated with rhIL-17 or CM of Th17 cells showed clear and long branch-like invadopodia demonstrating higher invasive potential of these cells and resulted in 1.5-fold (orange line) or 1.6-fold (blue line) higher invasion rate after 5 days. In summary, our results showed that Th17 cells induced an EMT phenotype in cervical cancer cells after direct cell-cell contact, however, stimulation with rhIL-17 or supernatants of Th17 cells was sufficient to induce the EMT phenotype and significantly promoted their migration and invasiveness.

### Th17 cells induced the expression of miR-142-5p in cervical cancer cells in an IL-17-dependent manner that mediated increased migration and invasiveness

Next, we were interested in the mechanism responsible for Th17-induced enhanced migration and invasion of cervical cancer cells. A microRNA expression analysis revealed up-regulation of miR-142-5p in cervical cancer cells after co-cultures with Th17 cells, rhIL-17 and CM Th17 cell stimulation (data not shown). We validated these findings by qPCR analysis in cervical cancer cells resulting in up to 3-fold increased expression of miR-142-5p after co-cultures with Th17 cells (Supplementary Fig. S1D), 2.5-fold increase after rhIL-17 stimulation (orange bars) and strong 4.6-17.9-fold enhanced expression after CM Th17 cell stimulation (blue bars) (Fig. 2A). Neutralization of IL-17 in the CM of Th17 cells significantly reduced miR-142-5p expression (Fig. 2B; 45.5-75.6 % reduction) clearly demonstrating that induction of miR-142-5p was dependent on Th17-derived IL-17. Over-expression of miR-142-5p by transient transfection in different cervical cancer cells (Fig. 2C; green bars; 2.5-8.5-fold increase) mediated an up to 43 % increased closure of the gap resulting in a significant decrease of relative area in scratch assays (Fig. 2D) and up to 2.6-fold enhanced migration in transwell migration assays (Fig. 2E). Moreover, spheroids from miR-142-5p transfected cells exhibited significantly higher invasive behavior (green line) in comparison to empty vector transfected cells (dotted line) resulting in 1.5-fold increased invasion rate after 5 days (Fig. 2F). Thus, our results demonstrated that Th17 cells induced miR-142-5p expression via IL-17 that mediated miR-142-5p enhanced migration and invasion of cervical cancer cells.

**Figure 2:**
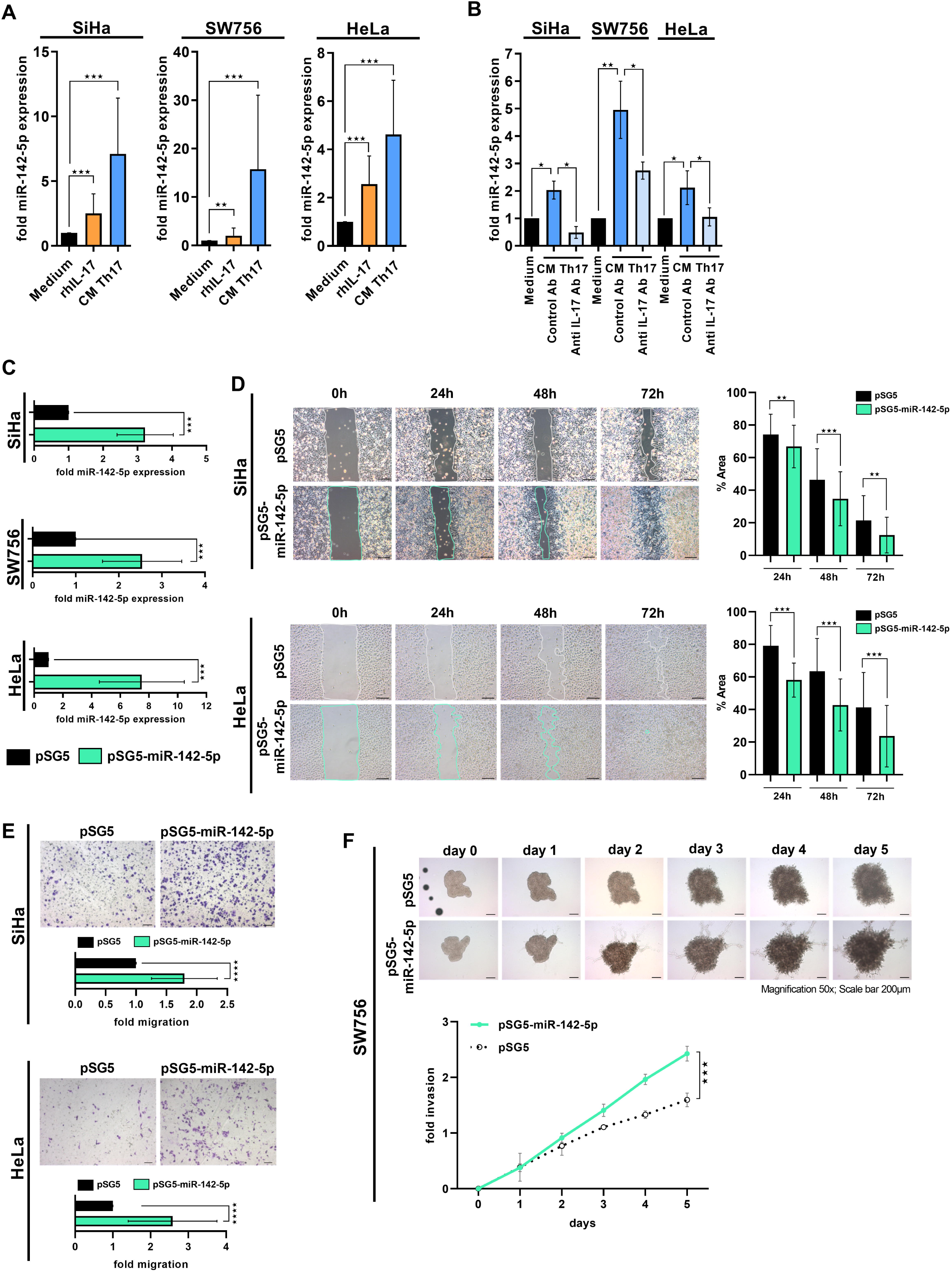
Th17 cells induced the expression of miR-142-5p in cervical cancer cells which mediated enhanced migration and invasion. (A) SiHa, SW756 and HeLa cells were stimulated with rhIL-17 (orange), CM of Th17 cells (blue) or medium (black bars) as a control for 24 h and miR-142-5p expression was analyzed in relation to RNU48. (B) In neutralization experiments, CM were prestimulated with neutralizing anti-IL-17 or respective isotype control antibodies for 2 h (light blue bars) and miR-142-5p expression was analyzed in relation to RNU48. (C) SiHa, SW756 and HeLa cells were transfected with miR-142-5p expressing plasmid (green bars) or pSG5 (black bars) as empty vector control. MiR-142-5p expression was analyzed in relation to RNU48 48 h later. (D) MiR-142-5p transfected SiHa and HeLa cells were scratched 24 h post-transfection. Pictures were taken after 0, 24, 48 and 72 h. The area of the gap, indicated by lines (left panel), was determined in relation to time point 0 h (right panel). Scale bar: 100 μm. Bars represents results (mean ±SD) of n=3 experiments. (E) MiR-142-5p transfected SiHa and HeLa cells were used in transwell migration assays 48 h post-transfection. Transmigrated cells were evaluated 24 h later. Numbers of pSG5 transfected cells were set at 1. Representative pictures (upper panel); quantification of n=3 experiments with five independent pictures, respectively (mean ±SD) lower panel. (F) Spheroids of mir-142-5p transfected SW756 cells were generated. Spheroids were embedded into matrigel, pictures were taken for 5 days and spheroid invasion was calculated. Shown are the results mean ± SD from three independent experiments performed in duplicates. P-value according to the nonparametric Kruskal-Wallis test (A, D) or Mann-Whitney U-test (B, C, E, F). Asterisks represent statistical significances: **P < 0.01; ***P < 0.001; ****P < 0.0001.

### Target prediction and validation of SDHC and SDHD as direct target genes of miR-142-5p by dual luciferase assay

To investigate how miR-142-5p contributes to enhanced migration and invasiveness of cervical cancer cells we performed an *in-silico* target prediction that identified subunits of the succinate dehydrogenase (SDH) complex with miR-142-5p binding sites in their 3′UTRs including SDHC and SDHD (Fig. 3A). We cloned the respective 3′UTR sequences of SDHC and SDHD into the pMIR-RNL-TK reporter vector and cotransfected these recombinants together with a miR-142-5p expression vector in HEK 293T cells. The luciferase activities (RLU (relative light units)) of the wild-type reporters were normalized against the RLU of the empty reporter vector. We found reduction of the RLU for SDHC (light blue bar, 21.3 % reduction, p=0.002) and SDHD (dark blue bar, 34.4 % reduction, p=0.0002) (Fig. 3B). To validate the direct binding of miR-142-5p to its target sites, we mutated the binding sites in the 3′UTRs of SDHC and SDHD. Both mutated reporter vectors (striped bars) showed a significant increase of the RLU when cotransfected with miR-142-5p compared with the respective non-mutated recombinants (18.9 % and 24.6 % increase, respectively) indicating the direct binding of miR-142-5p to its binding sites in the respective 3′UTRs (Fig. 3B). Analyzing the downstream effects of miR-142-5p over-expression on the endogenous protein level western blot analysis revealed significantly reduced expression of SDHC and SDHD in miR-142-5p transfected SiHa, SW756 and HeLa cells (23.1-60 % reduction, Fig. 3C, lower pannels). Thus, our results identified und validated SDHC and SDHD as new targets for miR-142-5p.

**Figure 3:**
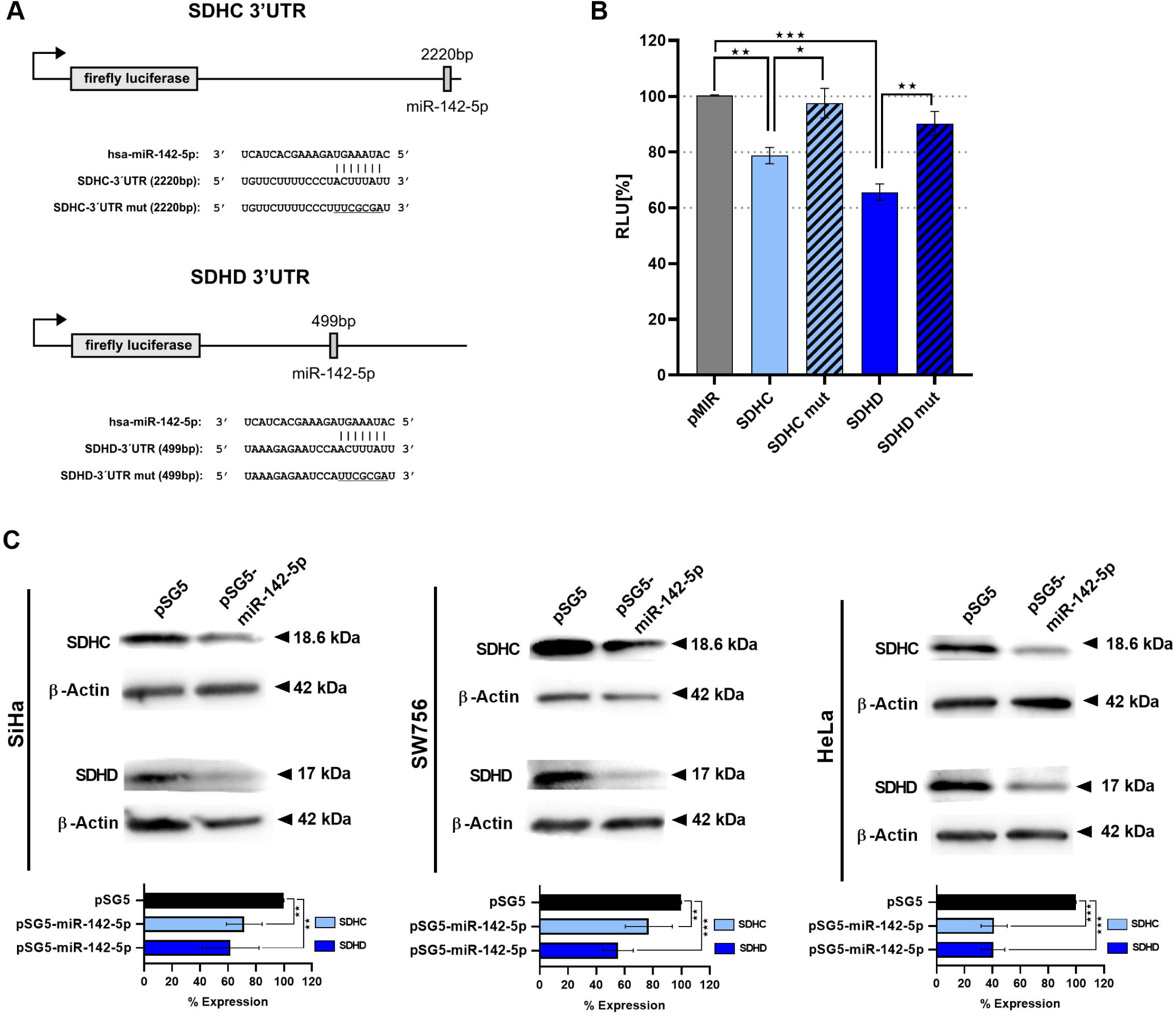
Identification and validation of SDHC and SDHD as new targets of miR-142-5p. (A) Schematic diagram of reporter gene plasmids. The position of the predicted miR-142-5p binding sites in the respective 3′UTR reporter plasmids and their corresponding sequences as well as the sequences of the mutated binding sites (underlined) are shown. SDHC-3’UTR reporter construct (upper panel), SDHD-3’UTR reporter construct (lower panel). (B) HEK 293T cells were transfected with the wild type reporter plasmids of the respective target genes SDHC (light blue bar) or SDHD (dark blue bar) or mutated reporter plasmids (mut) of the target genes (striped bars). The luciferase activities were normalized with respect to the luciferase activity measured with empty reporter construct (grey bar). The results represent the mean of three independent experiments carried out in duplicates. (C) SiHa, SW756 and HeLa cells were transfected with miR-142-5p expressing plasmid or pSG5 as empty vector control. Whole cell extracts were analyzed 72 h later for SDHC (light blue bars) or SDHD (dark blue bars) expression by western blot analysis. β-actin was used as a loading control. Bars represent quantification of n=3 independent experiments (mean ±SD). P-value according to the nonparametric Mann-Whitney U-test. Asterisks represent statistical significances: *P < 0.05; **P < 0.01; ***P < 0.001.

### Th17 cells reduced the expression of SDH subunits C and D and SDH activity in cervical cancer cells

Next, we analyzed the impact of Th17 cells on the expression of SDHC and SDHD in cervical cancer cells. In mRNA expression analysis based on microarrays, co-cultures between Th17 cells and cervical cancer cells (fold change of 0.86-0.84) and to a higher extend stimulation with rhIL-17 and CM of Th17 cells decreased the expression of SDHC (fold change of 0.78-0.60) and SDHD (fold change of 0.72-0.52) in SiHa and SW756 cells (data not shown). QPCR analysis revealed diminished SDHC and SDHD expression in cervical cancer cells after direct cell-cell contact with Th17 cells (Supplementary Fig. S2A; 20-52 % reduction) and stronger reduced expression of SDHC (31-50 % reduction, light blue bars) and SDHD expression (65-82 % reduction, dark blue bars) in three different cervical cancer cells after stimulation with CM of Th17 cells (Fig. 4A).

**Figure 4:**
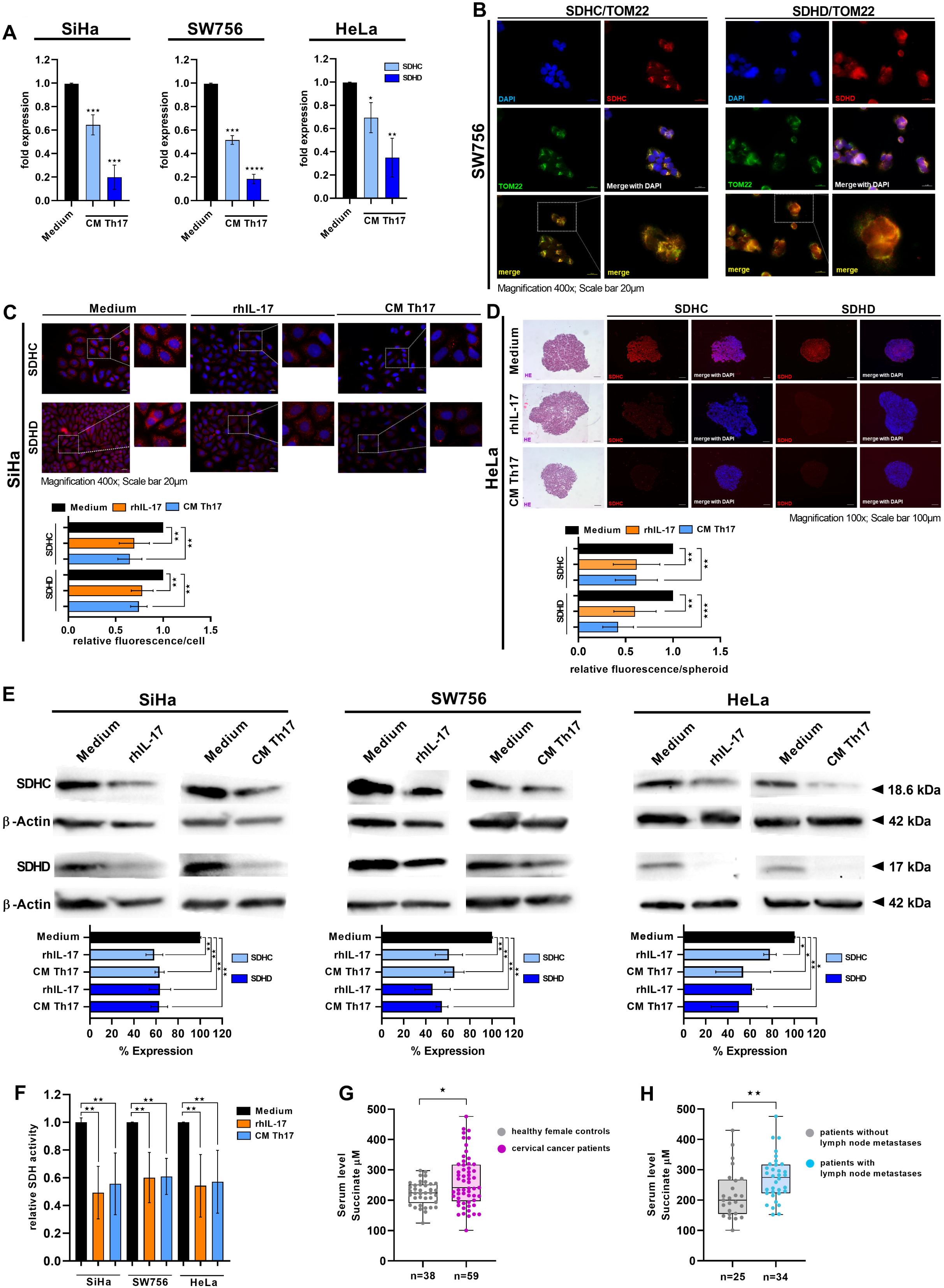
Th17 cells reduced the expression of SDHC and SDHD in cervical cancer cells. (A) SiHa, HeLa and SW756 cells were stimulated with CM of Th17 cells or medium for 48 h. Expression of SDHC and SDHD was evaluated by qRT-PCR. (B) SW756 cells were analyzed for SDHC, SDHD and TOM22 expression by IF. (C) 2D monolayers of SiHa cells were stimulated with rhIL-17, CM of Th17 cells or medium for 72 h and SDHC and SDHD expression was investigated by IF. Bars represent quantification of relative fluorescence/cell of 20 independent pictures (magnification 400x) from n=2 independent experiments performed in duplicates. (D) 3D spheroids of HeLa cells were generated over 10 days in the presence of medium, rhIL-17 or CM of Th17 cells. 5 µm sections of fixed paraffin-embedded spheroids were validated by HE stainings and analyzed for SDHC and SDHD expression by IF. Bars represent quantification of relative fluorescence/spheroid of n=6 independent spheroids, respectively. (E, F) SiHa, SW756 and HeLa cells were stimulated with medium, rhIL-17 or CM of Th17 cells for 72 h. (E) Whole cell extracts were analyzed for SDHC and SDHD expression by western blot analysis. β-actin was used as a loading control. Bars represent quantification of n=3 independent experiments (mean ±SD; SDHC: light blue bars; SDHD: dark blue bars). (F) SDH activity of SiHa, SW756 and HeLa cells stimulated with medium (black bars), rhIL-17 (orange bars) or CM of Th17 cells (blue bars) was determined after 72 h. Shown are the results from n=3 independent experiments (mean ±SD) performed in triplicates. P-value according to the nonparametric Kruskal-Wallis test. Asterisks represent statistical significances: *P < 0.05; **P < 0.01; ***P < 0.001; ****P < 0.0001. (G) Serum samples of 59 cervical cancer patients (purple dots) and 38 healthy female controls were analyzed for succinate. (H) Serum levels of succinate were evaluated in patients with (n=34; blue dots) and without (n=25; grey dots) lymph node metastases. Black line: median value of the respective groups. P-value according to the nonparametric Mann-Whitney U-test. Asterisks represent statistical significances: *P < 0.05; **P < 0.01.

To validate the results on protein levels we first analyzed SDHC and SDHD expression by IF (Fig. 4B, C). Co-stainings with TOM22 (green) confirmed co-localization of SDHC and SDHD expression (red) with the mitochondrial marker (Fig. 4B). We found significantly reduced expression of SDHC and SDHD after stimulation with rhIL-17 and CM of Th17 cells (33-45 % reduction in relative fluorescence/cell) in SiHa cells (Fig. 4C) and SW756 cells (Supplementary Fig. S3A). We further evaluated spheroids of cervical cancer cells showing increased invasiveness after stimulation with rhIL-17 and CM of Th17 cells (Fig. 1E). Sections of those spheroids showed reduced SDHC and SDHD expression in IF (Fig. 4D; 39-58 % reduction and Supplementary Fig. S3B) as well as, to a minor degree, Th17-cervical cancer cells heterospheroids (Supplementary Fig. S2B, C; 50-57% reduction). Furthermore, western blot analysis revealed significant reduction of SDHC (light blue bars) and SDHD (blue bars) after stimulation with rhIL-17 and CM of Th17 cells in three different cell lines (Fig. 4E). Finally, rhIL-17 (orange bars) as well as CM of Th17 cells (light blue bars) significantly diminished activity of the SDH complex in all three tested cervical cancer cells (Fig. 4F; 40-51 % reduction). In summary, we could show that Th17 cells reduced the expression of SDHC and SDHD as well as activity of the SDH complex in cervical cancer cells.

As the SDH complex catalyzes the oxidation of succinate to fumarate linking the TCA cycle to oxidative phosphorylation (17) and reduced SDH complex activity is associated with succinate accumulation, we evaluated serum samples from 59 cervical cancer patients to validate our *in vitro* findings *in vivo*. We found enhanced concentration of succinate in the patients serum′ (purple dots) in comparison to 38 female healthy age-matched controls (grey dots; Figure 4G). Notably, patients with lymph node metastases exhibited significantly higher levels of succinate in their serum (blue dots; 1.4-fold increased) in comparison to patients without metastases (grey dots; Figure 4H).

### Th17-induced migration of cervical cancer cells is dependent on miR-142-5p-mediated suppression of SDHC and SDHD

To analyze the relevance of SDHC and SDHD for migration of cervical cancer cells we knocked down SDHC or SDHD with two specific siRNAs in three cervical cancer cells (Fig. 5A). Besides the reduction of SDHC and SDHD expression (Fig. 5A), these knock downs affected EMT marker expression (Supplementary Fig. S4A, B) and led to reduced e-cadherin and enhanced vimentin protein expression (Supplementary Fig. S4C, D) supporting an EMT phenotype. In scratch assays with SiHa cells, the gap was closed significantly faster by cells in which SDHC (light blue dotted and striped bars), SDHD (blue dotted and striped bars) or both together (purple bars) were specifically knocked down (Fig. 5B). Furthermore, knock down of SDHC and SDHD increased migration of SiHa and HeLa cells (Fig. 5C, 1.6-2.4-fold increase) in transwell migration assays.

**Figure 5:**
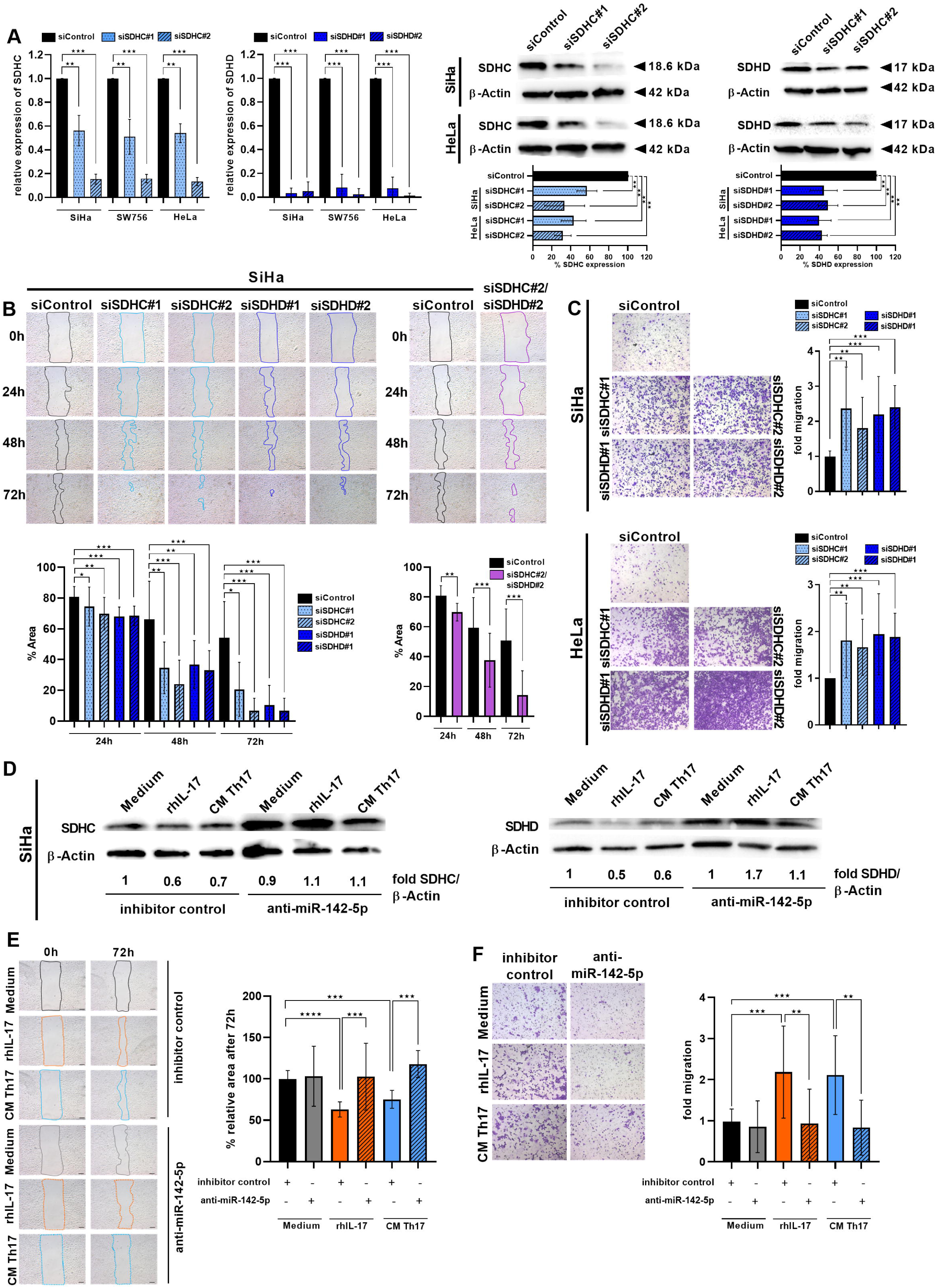
Th17-induced increased migration of cervical cancer cells is dependent on miR-142-5p-mediated reduced SDHC and SDHD expression. SiHa, SW756 and HeLa cells were transfected with two specific siRNAs for SDHC (dotted and stripped light blue bars) or SDHD (dotted and stripped dark blue bars), respectively, or mock siRNA (black bars) as a control. (A) After 48 h, SDHC and SDHD expression was calculated by qRT-PCR (left panel) and whole cell extracts were analyzed for SDHC and SDHD expression by western blot analysis (right panel). β-actin was used as a loading control. Bars represent quantification of n=3 independent experiments performed in duplicates. (mean ±SD). (B) 24 h post-transfection, SiHa cells were scratched. Pictures were taken after 0, 24, 48 and 72 h. The area of the gap, indicated by lines (upper panel), was determined in relation to time point 0 h (lower panel). Scale bar: 100 μm. Bars represent results (mean ±SD) of n=3 experiments. Additionally, SiHa cells were transfected with specific siRNAs for SDHC and SDHD (purple bars). 24 h post-transfection, SiHa cell were scratched. Pictures were taken after 0, 24, 48 and 72 h. The area of the gap, indicated by lines (upper panel), was determined in relation to time point 0 h (lower panel). Scale bar: 100 μm. Bars represents results (mean ±SD) of n=3 experiments. (C) Transfected SiHa and HeLa cells were used in transwell migration assays 48 h post-transfection. Representative pictures (left panel); quantification of transmigrated cells of n=3 experiments with five independent pictures, respectively (mean ±SD, right panel). (D, E, F) SiHa cells were transfected with inhibitor of miR-142-5p or inhibitor control. After 24 h, cells were stimulated with medium, rhIL-17 or CM of Th17 cells. (D) Whole cell extracts were analyzed for SDHC and SDHD expression by western blot analysis. β-actin was used as a loading control. Fold expression of SDHC and SDHD per β-actin expression was calculated. Medium stimulated cells transfected with inhibitor control was set at 1. (E) Transfected cells were stimulated with medium, rhIL-17 (orange bars) or CM of Th17 cells (light blue bars). After 24 h cells were scratched. Pictures were taken after 0, 24, 48 and 72 h. The area of the gap, indicated by lines (left panel), was determined after 72 h in relation to time point 0 h (right panel). Scale bar: 100 μm. Bars represent results (mean ±SD) after 72 h of n=3 experiments. The area of medium stimulated cells transfected with inhibitor control was set at 100 %. (F) 24 h post-transfection, transfected cells were stimulated with medium, rhIL-17 (orange bars) or CM of Th17 cells (light blue bars). After 24 h, cells were used in transwell migration assays. Representative pictures (left panel) after 24 h; quantification of transmigrated cells of n=3 experiments with five independent pictures, respectively (mean ±SD; right panel). The number of medium stimulated cells transfected with inhibitor control was set at 1. P-value according to the nonparametric Kruskal-Wallis test. Asterisks represent statistical significances: **P < 0.01; ***P < 0.001; ****P < 0.0001.

To validate the role of miR-142-5p-mediated down-regulation of SDHC and SDHD for Th17-induced migration and invasion of cervical cancer cells, we transfected cervical cancer cells with inhibitor of miR-142-5p or an inhibitor control, stimulated the cells with rhIL-17, CM of Th17 cells or medium and analyzed their proliferation and migration (Fig. 5D, E, F). Transfection with inhibitor of miR-142-5p reverted Th17-mediated suppression of SDHC and SDHD (Fig. 5D) resulting in 40-90 % increased SDHC and SDHD expression. Stimulation of inhibitor control transfected cells with rhIL-17 (orange bars) or CM of Th17 cells (light blue bars) resulted in faster closure of the gap in scratch assays (Fig. 5E) and enhanced migration in transwell migration assays (Fig. 5F). In contrast, transfection of inhibitor of miR-142-5p significantly reduced proliferation and migration of IL-17-(striped orange bars) and Th17-instructed (striped blue bars) cervical cancer cells (Fig. 5E, F). Taken together, our data clearly showed that Th17-induced enhanced migration of cervical cancer cells is dependent on miR-142-5p-mediated suppression of SDHC and SDHD.

### Th17 cells correlate with diminished SDHD expression in cervical cancers *in situ* associated with lymph node metastases and recurrent cervical cancers

To investigate the relationship between Th17 cells and SDH expression in cervical cancer patients, we analyzed SDHD expression by IHC in 52 clinical biopsies (Fig. 6A). In our experiments, SDHD showed strongest reduction in expression within the SDH subunits mediated by Th17 cells (Fig. 4A). Applying the IRS, we found a heterogeneous SDHD expression *in situ* reaching from negative to strong (representative pictures in Fig. 6A). 12/52 were negative for SDHD expression in cervical cancer cells exhibiting partial stromal SDHD expression. 40/52 were judged as positive with weak (n=23), moderate (n=15) or strong (n=2) SDHD expression (Fig. 6B). We found a decline in SDHD expression during cervical cancer progression based on tumor FIGO stages resulting in a weak or absent SDHD expression in more advanced cervical cancers (Fig. 6C; r=-0.6228; p<0.0001). Interestingly, patients with lymph node metastases showed significantly less SDHD expression in their tumor tissues in comparison to patients without metastases (Fig. 6D). Moreover, retrospective analysis demonstrated that in patients developing recurrence of cervical cancers SDHD expression was significantly less frequent in their tumor tissues than in patients without relapse (Fig. 6E). When we analyzed the relation between the presence of Th17 cells and SDHD expression we found a significant inverse correlation between the numbers of tumor-infiltrating Th17 cells/mm^2^ and IRS of SDHD (Fig. 6F; r=-0.7073) whereas no correlation was found between numbers of CD4^-^ IL-17^+^ cells /mm2 and SDHD expression (Supplementary Fig. S5A). The correlation was intensified when we evaluated the proportion of Th17 cells per total infiltrating CD4^+^ cells per biopsies (as percentages of CD4^+^IL-17^+^ cells/CD4^+^ cells) (Fig. 6G; r=-0.7833). Thus, our results indicated a functional link between Th17 cells and reduced SDHD expression during cervical cancer progression *in vivo*.

**Figure 6:**
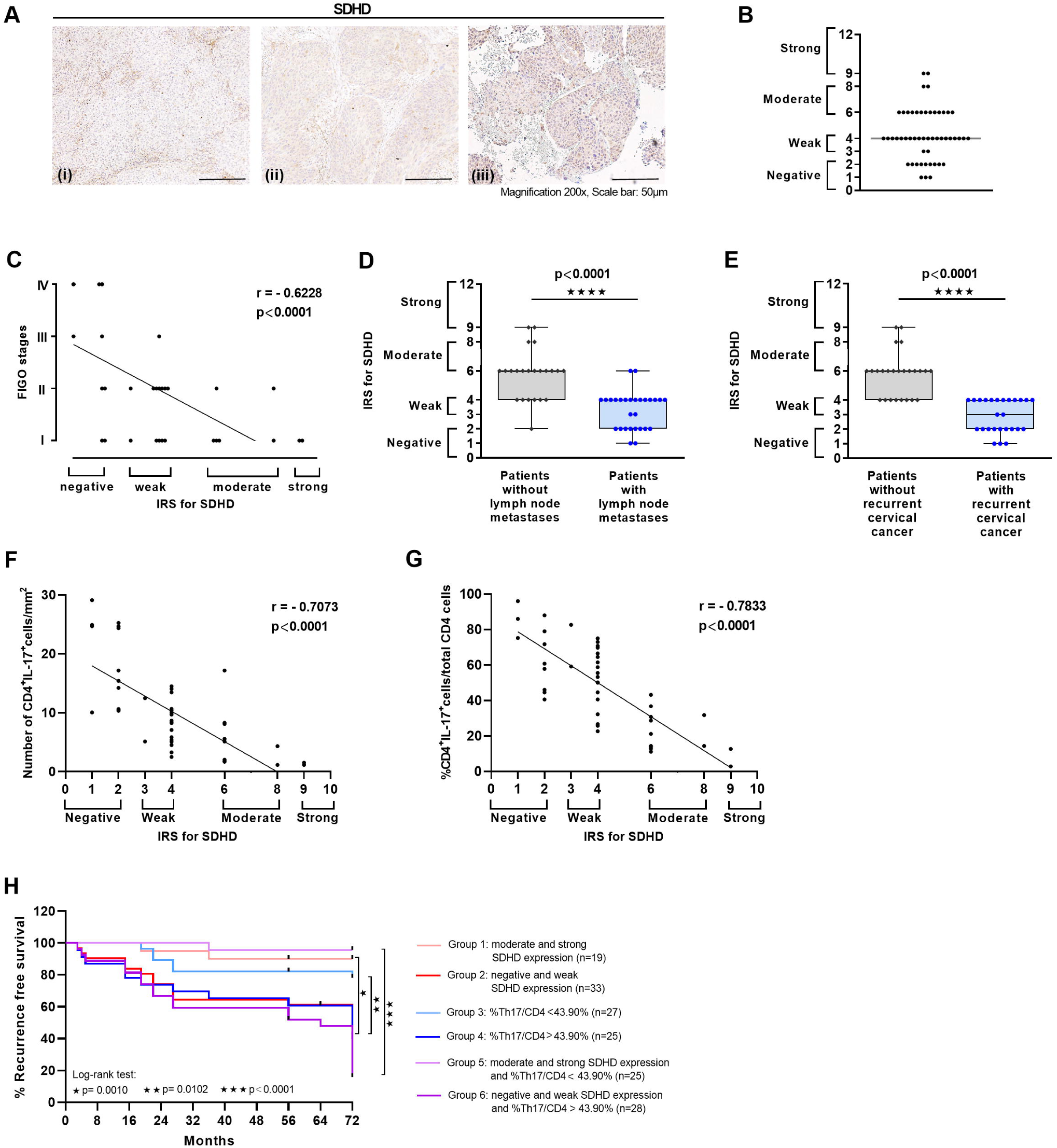
Inverse correlation of Th17 cells and SDHD expression in cervical cancer biopsies *in situ* and association with lymph node metastases and cervical cancer recurrence. Sections of human SCCs of 52 cervical cancer patients were stained for SDHD expression by IHC ((i) negative, (ii) weak, (iii) moderate expression). Magnification 200x (Scale bars 50 μm). (B) IRS of SDHD within 52 biopsies. (C) Correlation of IRS of SDHD and tumor FIGO stages. (D) IRS of SDHD in tumor tissues of patients with (blue circles) and without (grey diamonds) lymph node metastases or (E) with and without cervical cancer relapse. (F, G) Correlation of numbers of Th17 cells/mm^2^ or (G) percentages of Th17/CD4^+^ T cells with IRS of SDHD. (H) Recurrence-free survival of 52 patients was determined for a cohort with moderate and strong SDHD expression (n=19; light red line) and negative and weak SDHD expression (n=33; red line). Median recurrence-free survival was 72 months for the cohort with negative and weak SDHD expression. Comparison of survival analysis was performed using log-rank (Mantel-Cox) test; chi-square: 10.83, P = 0.0010. Recurrence-free survival of 52 patients was determined for a cohort with CD4^+^IL-17^+^ cells per CD4^+^ T cells < 43.90 % (n=26; light blue line) and CD4^+^IL-17^+^ cells per CD4^+^ T cells > 43.90 % (n=24; blue line). Median recurrence-free survival was 72 months for the cohort with CD4^+^IL-17^+^ cells per CD4^+^ T cells 43.90 %. Comparison of survival analysis was performed using log-rank (Mantel-Cox) test; chi-square: 6.61, P = 0.0102. Recurrence-free survival of 52 patients was determined for a cohort with moderate and strong SDHD expression and CD4^+^IL-17^+^ cells per CD4^+^ T cells < 43.90 % (n=23; light purple line) and for a cohort with negative or weak SDHD expression and CD4^+^IL-17^+^ cells per CD4^+^ T cells > 43.90 % (n=27; purple line). Median recurrence-free survival was 64 months for the cohort with negative or weak SDHD expression and CD4^+^IL-17^+^ cells per CD4^+^ T cells > 43.90 %. Comparison of survival analysis was performed using log-rank (Mantel-Cox) test; chi-square: 31.61, P < 0.0001.

To evaluate whether the presence of Th17 cells and reduced SDHD expression is linked to course of disease, ROC analyses were performed to identify a cutoff value concerning SDHD expression and Th17 frequencies which discriminates between patients with and without recurrent cervical cancers. Best discrimination was obtained for a cut-off value of Th17 per CD4^+^ T cells > 43.9 % (100 % sensitivity and 93.3 % specificity; Supplementary Fig. S5B) and an IRS score < 6 (weak or negative SDHD expression; 100 % sensitivity and 65.4 % specificity; Supplementary Fig. S5C)). The area under the ROC curve (AUC) value was 0.9201 and 0.9956, respectively. Applying the defined cut-off values for 52 patients (Fig. 6H), in the group of patients with moderate and strong SDHD expression (light red line) RFS was 90 % after 6 years, while for patients with negative or weak SDHD expression (red line) 3-and 6-years RFS was 60.8 and 42.1 %, respectively. Regarding Th17 frequencies, for patients with percentages of Th17 per CD4^+^ T cells < 43.9 % (light blue line) 6-years RFS was 78.3 % and for patients with percentages of Th17 per CD4^+^ T cells > 43.9 % (blue line) 6-years RFS was 42.1 %. Interestingly, when we combined both factors, Th17 frequencies and SDHD expression, in the group of patients with moderate and strong SDHD expression and %Th17/CD4^+^ T cells < 43.9 % (light purple line) 6-years RFS was 96 %. In contrast, in the group of patients with weak and negative SDHD expression and percentages of Th17 per CD4^+^ T cells > 43.9 % (purple line), 3-and 6-year recurrence-free survival was 59.3 % and 15.6 %, respectively. In conclusion, our data demonstrated a clear association between Th17 levels as well as reduced SDHD expression and cervical cancer recurrence.

## Discussion

Th17 cells infiltrate cervical cancers and are associated with poor prognosis for the patients (6, 10). In this study, we investigated how Th17 cells support cervical cancer progression favoring metastases and relapse. We could show that Th17 cells promoted migration and invasive behavior of cervical cancer cells that was dependent on Th17-induced IL-17-dependent miR-142-5p expression. We identified the subunits C and D of the succinate dehydrogenase complex as new targets of miR-142-5p mediating enhanced invasiveness of cervical cancer cells. Notably, patients with lymph node metastases exhibited high serum levels of succinate and *in situ* analysis revealed a correlation between numbers of Th17 cells and reduced SDHD expression that was associated with cervical cancer relapse. Fig. 7 summarizes our current concept of a novel Th17-induced mechanism favoring cervical cancer progression.

**Figure 7:**
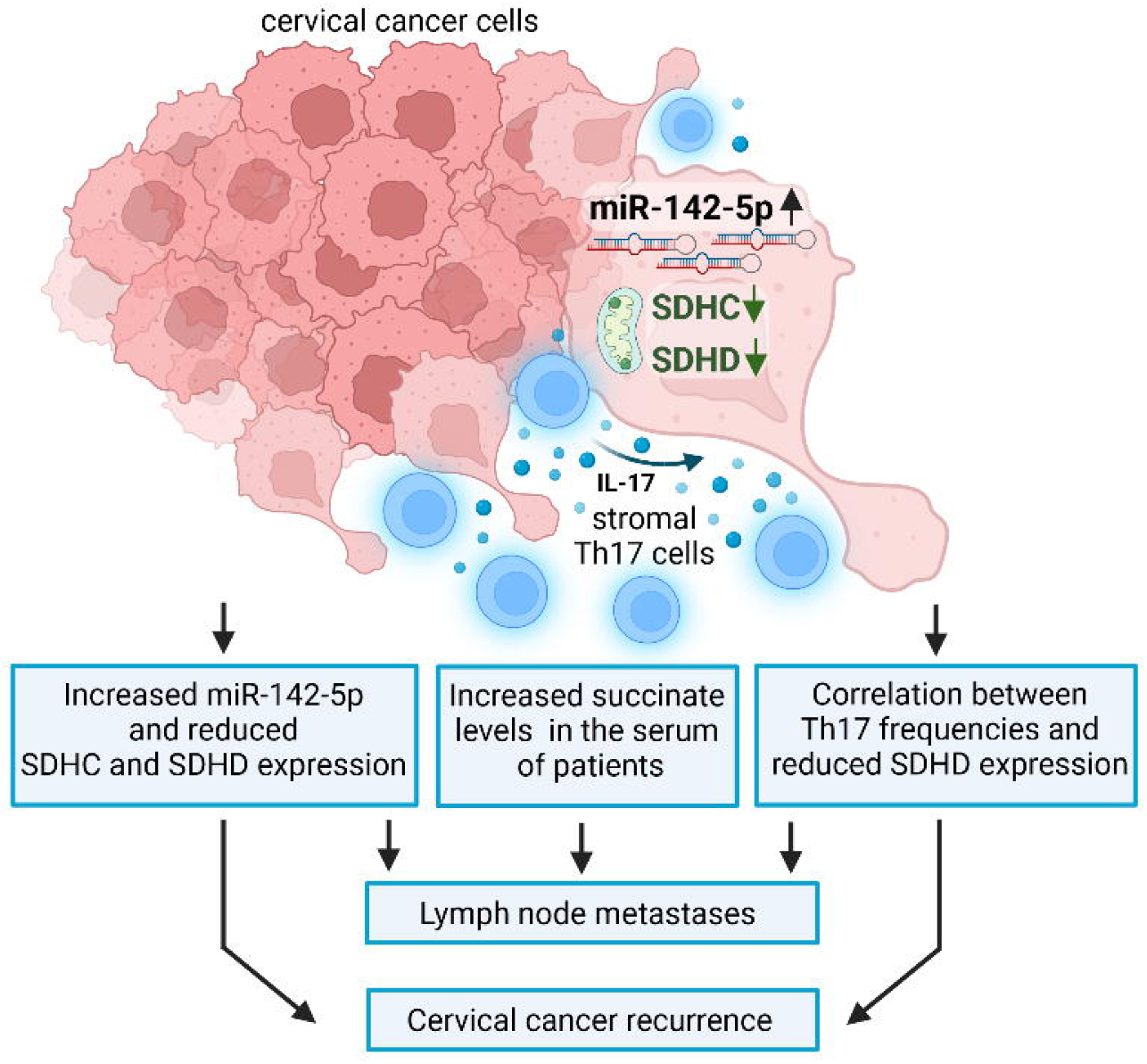
Schematic presentation of the role of Th17 cells in cervical cancer progression: Th17-induced miR-142-5p-dependent reduction in SDHC and SDHD expression favors cervical cancer invasiveness and increased frequencies of Th17 cells and reduced SDHD expression related to cervical cancer recurrence (scheme was generated with the BioRender software).

Metastatic cervical cancer remains a major clinical challenge. Prognosis of metastatic cervical cancer patients remains poor with a median survival time of only 8-13 months and a 5-year survival rate of 16.5 % (23, 24). The constitution of HPV-associated individual tumor microenvironment affects tumor progression either positively correlated with clinical outcome (25) or favoring a tumor-promoting environment (5). Our previous *in situ* analysis linked the presence of Th17 cells with cancer progression (11), reduced response toward chemoradiotherapy (6), lymph node metastases and cervical cancer recurrence (10). In this study, we showed that Th17 cells favored an EMT phenotype of cervical cancer cells. This occurred after direct cell-cell contact in co-cultures between Th17 cells and cervical cancer cells, however, Th17-produced soluble factors were sufficient to induce the effect without direct cell-cell contact and increased migration and invasiveness of cervical cancer cells. Th17 cells have a controversial role in different cancers. They mediate antitumor effects by recruiting immune cells into tumors or stimulating effector CD8^+^ T cells (26) as well as tumor-promoting responses driving proliferation, invasion, metastasis, and angiogenesis (27, 28). Although the presence of Th17 cells *in situ* was linked with cervical cancer progression (6, 10), and IL-17A was found to upregulate MMP-2 and MMP-9 expression favoring migration of cervical cancer cell lines (27), Th17-induced mechanisms underlying cervical cancer metastases are not reported so far.

Besides the infection with high risk HPV as the primary risk factor for the development of cervical cancers (1), other factors were considered to promote cervical cancer progression, like microRNAs (29). These highly-conserved 19 to 24 nucleotides long double-stranded RNAs regulate gene expression involved in carcinogenesis favoring resistance, invasion and metastases (30). During cervical carcinogenesis, aberrant microRNA expression profiles were described and several microRNAs were identified as OncomiRs promoting cervical cancer progression (29). Different studies found an increased expression of miR-142-5p in cervical cancers in comparison to normal cervical tissues that favored cervical cancer progression by targeting Wnt/β-catenin pathway (31) and SMAD3, a negative regulator of TGF-β signaling (32). In other cancer entities, increased levels of miR-142-5p were associated with progression of colon (33), breast (34), skin (35) and renal cancer (36). Here we could show that Th17 cells, representing a part of the inflamed tumor-promoting cervical cancer micromilieu, increased the miR-142-5p expression in cervical cancer cells, on the one hand after direct cell-cell contact by co-cultures, on the other hand stronger by soluble factors that were sufficient for miR-142-5p induction enhancing migration and invasiveness of 2D cultures and 3D spheroids. Targeting of Th17-derived IL-17 via neutralizing antibodies identified IL-17 as the responsible inducer of miR-142-5p. This is in line with our previous findings identifying Th17-produced IL-17 as the responsible soluble factor that mediate resistance of cervical cancer cells toward chemoradiotherapy (6). Thus, our results identified a second tumor-promoting mechanism mediated by IL-17 promoting cervical cancer progression.

In search for a responsible mediator of Th17-induced miR-142-5p-driven enhanced invasiveness of cervical cancer cells, we identified the subunits C and D of the SDH complex whose expression and activity were reduced by rhIL-17 or CM of Th17 cells and which may act in concert with other described miR-142-5p targets favoring cancer progression. The SDH complex assumes a prominent role in different cancers whose development is linked to mutations within the SDH complex (18, 19) or which show reduced expression of individual SDH subunits promoting EMT, aerobic glycolysis and Warburg effect (20, 21). In line with this, our knock down of SDHC or SDHD in cervical cancer cells increased the expression of vimentin, ZEB1 and TWIST, but reduced e-cadherin. Mechanistically, the EMT phenotype in ovarian cancer cells after knock down of the subunit B of the SDH complex was linked to histone hypermethylation (37). Furthermore, a feed-forward loop initiated by SDHC knock-out in breast cancer cells mediated enhanced TWIST1 and SNAI1 expression favoring SDHC suppression and reduction of SDH complex activity (19). Additionally, besides this direct impact of the SDH subunits on EMT phenotypes, another study claimed their indirect role mediated via subsequent succinate accumulation that favors EMT and cancer cell migration via Succinate Receptor 1 (SUCNR1) (38). In our study, we identified that Th17 cells decreased the expression of SDHC and SDHD in cervical cancer cells resulting in diminished SDH complex activity that was mediated by miR-142-5p and validated SDHC and SDHD as new targets of miR-142-5p. Together with results of a previous study, describing SDHB as a target of miR-142-5p in colon cancer (36), our data indicated that miR-142-5p, and thereby Th17 cells, targets three of four subunits of the SDH complex and consequently reduces its activity. This may have major implications on cervical cancer metabolism favoring the accumulation of succinate, a defined oncometabolite (39). Consistently, we found enhanced succinate levels in the serum of cervical cancer patients in comparison to healthy female controls and highest succinate levels were linked with lymph node metastases suggesting evaluation of a potential role of succinate for cervical cancer progression in further studies. Thus, the findings from this study unraveled a novel molecular mechanism favoring cervical cancer progression and may contribute to a better understanding of the link between the presence of Th17 cells and course of disease with poor clinical outcome for cervical cancer patients.

Clinically most important, retrospective *in situ* analysis from this study further supported the *in vivo* relevance of our findings. When we evaluated the presence of SDHD in cervical cancer biopsies, the SDH subunit with strongest reduction in expression mediated by Th17 cells, we found a heterogeneous expression of SDHD within the biopsies and diminished expression of SDHD correlated with advanced tumor FIGO stages, lymph node metastases and cervical cancer recurrence. Interestingly, enhanced proportion of Th17 cells correlated with reduced SDHD expression *in situ* supporting our *in vitro* findings. IL-17 expression was not solely found in T cells in cervical cancers (6, 40); however, our analysis revealed that numbers of IL-17-expressing CD4 negative cells are not linked to SDHD expression in cervical cancers. Both parameters, negative or weak SDHD expression and a ratio of Th17/CD4^+^ T cells > 43.90%, were associated with reduced recurrence free survival. Notably, after using a combination of both parameters, the 3-and 6-year recurrence-free survival was 59.3 % and only 15.6 %. In conclusion, numbers of Th17 cells increase (10, 11), while SDHD expression decreases during cervical carcinogenesis (this study) and both are associated with severity of the disease.

For patients who do not respond to standard treatments, that favor metastases and relapse, new strategies like new immunotherapeutic approaches are needed. So far, two antibodies are approved as targeted immunotherapies in combination with a platinum-based chemotherapy in patients with advanced, metastatic or recurrent cervical cancer: bevacizumab, an anti-VEGF antibody to inhibit angiogenesis (41), and pembrolizumab, an anti-PD1 antibody (42). Our results from this study highlighted the relevance of the Th17-miR-142-5p-SDH axis for cervical cancer invasiveness suggesting this axis as potential biomarker or target for immunotherapy. MicroRNAs are currently being discussed as an attractive target for anticancer drug development (43). Moreover, the SDH complex is considered as potential therapy target on molecular level inducing SDH activity by SDH activators (44). On the basis of our results, Th17 numbers *in situ* as initiators of the novel mechanism should be evaluated as target for immunotherapy. For the treatment of psoriasis, antibodies against IL-17A or IL-17 receptor are approved (45) and are currently being evaluated for treatment of inflammatory diseases (46) and cancers (47). In line with this, our results identified IL-17 as the responsible initiator of the tumor-promoting miR-142-5p-SDH-axis further supporting the idea of Th17-based immunotherapeutic approaches in cervical cancers.

In conclusion, our study identified a novel role of Th17 cells in cervical cancer progression and metastases. We could clarify that Th17 cells induced miR-142-5p affecting activity of the SDH complex in cervical cancer cells and thereby supported cancer migration and invasiveness. Our results may explain the association of Th17 cells in cervical cancers and cancer progression favoring the idea of Th17-based immunotherapeutic approaches in cancer treatments.

## Material and methods

### Cell culture

HPV18-positive cervical carcinoma cell lines SW756 (RRID:CVCL_1727), HeLa (RRID:CVCL_0030) and HPV16-positive SiHa cells (RRID:CVCL_0032) were received from ATCC or DSMZ and authenticated by a multiplex human cell line authentication test (Multiplexion, DSMZ) in August, 2020. Cells were cultured in DMEM (Sigma-Aldrich, Taufkirchen, Germany) supplemented with 10 % heat-inactivated endotoxin-tested fetal calf serum (FCS) (Biochrom, Berlin, Germany), 1 mM sodium pyruvate, and 2 mM L-alanyl-L-glutamine. Mycoplasma testing was performed monthly using Venor®GeM Classic (Minerva Biolabs; Berlin, Germany).

### Generation and cultivation of Th17 cells *in vitro* and ELISA

Naive CD4-positive T cells were isolated by negative selection from freshly obtained PBMC using the Human Naive CD4+ T-Cell Isolation Kit II (Miltenyi Biotech, Bergisch-Gladbach, Germany). Cells were cultured in RPMI-1640 medium supplemented with 10 % heat-inactivated endotoxin-tested FCS (Biochrom) and stimulated with the T-cell Activation/Expansion Kit (Miltenyi Biotech) for 7 to 10 days in the presence of recombinant IL-1β (10 ng/mL), IL-6 (40 ng/mL), IL-23 (10 ng/mL), and TGFβ (5 ng/mL; Miltenyi Biotech) (6, 48). CD4^+^/IL17^+^ cells were isolated using stimulation and staining reagents from the IL17-Secretion Assay-Detection Kit (Miltenyi Biotech). Purity was determined with anti-CD4-PE (AB_395752) and anti-CCR6-AlexaFluor647 (AB_1645402), or isotype controls and analyzed by flow cytometry (FACSCanto II; BD Biosciences). For conditioned media (CM) Th17 cells were cultured at a density of 1 x 10^6^/ml. After 24 h, fresh RPMI1640 medium (Sigma) plus supplements (10 % heat-inactivated endotoxin-tested FCS (Biochrom) and 1mM sodium pyruvate) was added. Conditioned media were collected 24 h later and analyzed for IL-17A by DuoSet ELISA (R&D Systems, Minneapolis, USA). Concentrations of IL-17A in the used CM was 764 pg/ml. In neutralization experiments neutralizing anti-IL-17 antibody (AB_354463) or matched isotype control antibody (AB_354267) (1 μg/ml); R&D Systems) was added to CM 2 h before usage.

### MRNA microarray

For detection of mRNA expression changes in rhIL-17 and CM Th17 stimulated SiHa and SW756 cells or in cervical cancer cells after co-cultures with Th17 cells the Agilent Low Input, one-color, Quick Amp Labeling Kit and SurePrint G3 Human Gene Expression 8×60Kv3 Microarray (Cat. No. G4851C, Agilent Technologies) was used corresponding to the manufacturer’s protocols. In brief, 100 ng total RNA was reversed transcribed at 40°C for 2 hours using T7 Primer. Subsequently, the cRNA was labeled with Cyanine3-pCp using the T7 RNA polymerase for 2 hours at 40°C. The purification of the labeled cDNA was performed with RNeasy Mini kit (Qiagen) according to Agilent’s One-Color Gene Expression Microarray Protocol followed by the quantification using a NanoDrop2000 Spectrophotometer (Thermo Fisher Scientific). The hybridization of 600 ng labeled cRNA to the microarray slide was carried out at 65°C, 17 hours, 10 rpm in the SureHyb chambers (Agilent Technologies). After two washing steps the microarray slide was scanned on the Agilent Microarray Scanner G2565BA (Agilent Technologies). The raw fluorescence signals were analyzed with the Agilent AGW Feature Extraction software (V.10.7.1.1, Agilent Technologies). Normalization of the background corrected values was done with the Biological Significance analysis using GeneSpring (V.14.9, Agilent Technologies). In our analysis only mRNAs with RefSeqAccession number were included that were detected in all quadruplicate of the tested samples. The fold change of the mRNAs from rhIL-17 and CM Th17 stimulated cells as well as cervical cancer cells after co-cultures was calculated by normalization of the expression values to the mean expression values of the control samples.

### MiRNA expression plasmid, reporter constructs and automated dual luciferase reporter assay

The pSG5-miR-142-5p expression plasmid was described previously (49). The sequences of the SDHC and SDHD 3′UTRs were cloned via *Spe*I and *Sac*I restriction sites into the pMIR-RNL-TK vector, which was described in Beitzinger et al. (50). The 3′UTR sequences of SDHC mut and SDHD mut were generated by overlap extension PCR using specific primers (Supplementary Table S1) replacing all predicted hsa-miR-142-5p target sites with *Nru*I restriction sites.

For automated dual luciferase reporter assay 2-2.5 x 10^4^ HEK 293 T cells were seeded per 96-well by the liquid handling system epMotion 5075 (Eppendorf, Hamburg, Germany). HEK 293 T cells were transfected with 50 ng/well reporter vector with or without 3′UTR and 200 ng/well pSG5 empty vector or pSG5-miR-142-5p expression plasmid, respectively. 48 h post transfection, cells were lysed and the cell lysates were prepared according to manual of the Dual-Luciferase® Reporter Assay System (Promega, Madison, USA) and measured with the GlowMax navigator microplate luminometer (Promega, Madison, USA).

### qRT-PCR

RNA was isolated using Qiazol (Qiagen, Hilden, Germany), and cDNA was generated from 1 μg of RNA with SuperscriptII (Invitrogen, Carlsbad, CA, USA). PCR primers (Sigma Aldrich, Taufkirchen, Germany) and probes (Roche Universal Probe Library; Roche) were designed using the Probe Finder software version 2.53 (Roche). The 140-bp fragment of SDHC was detected with primers 5′-ATGGGATCCGACACTTGATG-3 and 5′-GCTGGGAGCCTCCTTTCTT-3 and probe no. 46; the 104-bp fragment of SDHD was detected with primers 5′-CTTGCTCTGCGATGGACTATT-3 and 5′-AAGCCCAGGAGTTGGGTAAT-3 and probe no. 71; the 94-bp fragment of RPL13A was detected with primers 5′-AGCGGATGAACACCAACC-3 and 5′-TTTGTGGGGCAGCATACTC-3 and probe no. 28; the 95-bp fragment of CDH1 was detected with primers 5′-GATGGCGGCATTGTAGGT-3 and 5′-GCTCTGTCATGGAAGGTGCT-3 and probe no. 5; the 113-bp fragment of VIM was detected with primers 5′-TGAGATTGCCACCTACAGGAA-3 and 5′-GAGGGAGTGAATCCAGATTAGTTT-3 and probe no. 11; the 129-bp fragment of TWIST1 was detected with primers 5′-GGCTCAGCTACGCCTTCTC-3 and 5′-CCTTCTCTGGAAACAATGACATCT-3 and probe no. 57 and the 105-bp fragment of ZEB1 was detected with primers 5′-GGAGGATGACACAGGAAAGG-3 and 5′-TCTGCATCTGACTCGCATTC-3 and probe no. 88. For miR-142-5p detection, cells were lysed using Qiazol and total RNA was isolated by miRNeasy Mini KIT (Qiagen). Total RNA (1µg) was reverse transcribed into cDNA by miScript RT II Kit using miScript HiSpec Buffer or miRCURY LNA RT Kit (Qiagen). Expression levels of miR-142-5p were analyzed using miScript PCR System or miRCURY LNA SYBR Green PCR Kit (Qiagen). Specific primer pairs (miScript Primer Assay or miRCURY LNA miRNA PCR Assay, Qiagen) for miR-142-5p and RNU48 were used. RNU48 served as endogenous control. Real-time PCR was performed with a StepOnePlus Real-Time PCR System (Applied Biosystems, Foster City, USA).

### Protein expression analysis by western blot and immunofluorescence (IF) and evaluation of SDH activity

Carcinoma cells were seeded at a density of 4×10^5^ (SiHa), 5×10^5^ (SW756) and 3×10^5^ (HeLa) cells/6-well culture dish. 24 h later, they were incubated with medium, rhIL-17 (100 ng/ml) or conditioned media of Th17 cells (diluted 1:5) for 72 h. To prepare whole cell extracts, cells were lysed with lysis buffer (130 mM Tris/HCl, 6 % SDS, 10 % 3-Mercapto-1,2-propanediol, 10 % glycerol) and 3 times treated with ultrasound for 5 s. 15 μg of the whole protein extracts was separated by SDS gel electrophoresis and transferred to a nitrocellulose membrane (Whatman, GE Healthcare, Freiburg, Germany). Rabbit polyclonal anti-SDHC (1:200; AB_2634976; Fisher Scientific), rabbit polyclonal anti-SDHD (1:200; AB_2717499; Fisher Scientific), mouse anti-e-cadherin (1:1000; AB_2728770; Cell Signaling), rabbit anti-vimentin (1:1000; AB_10695459; Cell Signaling) and mouse anti-β-Actin antibody (1:5000; AC-15; AB_476744; Sigma-Aldrich) were used. Secondary Abs (Sigma-Aldrich) and SuperSignalWest Dura Substrate (Thermo Fisher Scientific, Schwerte, Germany) were used for detection with ChemiDoc XRS+ Molecular Imager. Expression was quantified with the Image Lab software (both Bio-Rad, Feldkirchen, Germany). Uncropped original western blots are shown in Supplementary Fig. S6. For IF, 3×10^4^ HeLa and SW756 cells were grown on glass coverslips, fixed, and permeabilized as described (11). Rabbit polyclonal anti-SDHC (1:200; AB_2634976; Fisher Scientific) and rabbit polyclonal anti-SDHD (1:200; AB_2717499; Fisher Scientific) were used followed by Alexa Fluor® 546-conjugated secondary antibody (Invitrogen). For visualization, an Olympus AX70 microscope with cellSens software from Olympus was used. To evaluate SDH activity cells were stimulated as described above and SDH activity was analyzed with Succinate dehydrogenase Activity Colorimetric Assay Kit (Sigma-Aldrich) according to the manufacturers′ instructions.

### Transient transfections and siRNA transfections

1×10^4^ SiHa or 1×10^3^ Hela cells/6-well were seeded and transfected with pSG5-miR-142-5p expression vector or pSG5 empty vector as a control using PolyFect Transfection Reagent (Qiagen, Hilden, Germany). After 48 h, miR-142-5p expression was analyzed by qPCR. 30 pmol of indicated siRNAs with respective target sequences (ON-TARGETplus Nontargeting siRNA #2, ON-TARGETplus siRNA #5 and #8 for human SDHC and siRNA #7 and #8 for human SDHD, all from Horizon, Cambridge, United Kingdom) per 1×10^4^ SiHa and 1×10^3^ HeLa cells/6-well culture dish were transfected with Lipofectamine RNAi-Max (Invitrogen). Efficient knock down was confirmed by western blot analysis. Cells were stimulated as described above and used in scratch and transwell migration assays.

### Scratch assay

For scratch assay, 4×10^5^ SiHa and HeLa cells/6-well were seeded and stimulated with rhIL-17 (100ng/ml), CM of Th17 cells or medium as a control. After 24 h, monolayers were scratched, washed with PBS and again stimulated with rhIL-17 (100ng/ml), CM of Th17 cells or medium as a control. The scratches were analyzed at time points 0, 24, 48 and 72 h with Leica DMI600B microscope and LAS X software. Cells transfected with SDHC or SDHD siRNA (1×10^4^) or pSG5-miR-142-5p expression plasmid (SiHa 1×10^4^; HeLa 1×10^3^) were scratched 24 h after transfection. Area of the scratch was quantified with ImageJ software (RRID:SCR_003070). Area of the respective scratches at time point 0 was set at 100 %.

### Transwell migration assay

For transwell migration assays, SiHa, SW756 and HeLa cells were stimulated with rhIL-17 (100ng/ml), CM of Th17 cells (diluted 1:5) or medium for 24 h. Cells were resuspended in DMEM without FKS and 5×10^4^ (SiHa, SW756) and 4×10^4^ (HeLa) were seeded into 24-well Transwell chambers (insert 6.5-mm diameter, 8-μm pore size; Corning Costar Corp.), loaded with DMEM containing 10% FKS. After 24 h, cells were fixed in 70 % ethanol and stained with crystal violet. Cells were washed with aqua dest. and non-migrated cells were removed from transwells using cotton swabs. Number of migrated cells was documented by five independent pictures per condition using Leica DMI600B microscope and LAS X software.

### Spheroid formation, spheroid invasion assay and IF of spheroids

4×10^3^ SW756 or HeLa cells were cultured in medium according to (51) in 48-well plates coated with 1% low attached agarose (Sigma Aldrich, Taufkirchen, Germany). For heterospheroids 4×10^3^ SW756 and 2×10^3^ Th17 cells were used. On day 5, spheroids were embedded into Matrigel® Growth Factor Reduced (GFR) Basement Membrane Matrix (Corning Costar Corp). Spheroid invasion was monitored over 5 days with Leica DMI600B microscope and LAS X software. Invasiveness was calculated using ImageJ software subtracting the area of the spheroids on day 0 from areas of day 1 to 5, respectively. For IF, spheroids were generated for 14 days, fixed in 4 % paraformaldehyde and embedded in paraffin. Five-µm-thick sections were cut and stained with hematoxylin and eosin (HE) according to standard procedures. For IF, antigen retrieval was performed by heating the sections in TE buffer (10mM Tris, 1mM EDTA pH 9.0) at 95°C for 15 min and immersed in 3 % H_2_O_2_ in TBS (50 mM Tris-HCl, 150 mM NaCl, pH 7.6) to block endogenous peroxidase activity. To reduce non-specific staining, each section was treated with 2.5 % normal horse serum for 30min. Rabbit anti-SDHC or rabbit anti-SDHD antibody (1:200, AB_2634976 and 1:200, AB_2717499, Fisher Scientific, Germany) as well as mouse anti-CD4 monoclonal antibody 4B12 (1:1000, Leica Biosystems; AB_563560) and rabbit anti-IL-17 polyclonal antibody (1:500, Abcam; AB_1603584) were used. For immunofluorescence TSA^TM^ Kit #2, with HRP-Goat Anti-Mouse IgG and Alexa Fluor ^TM^ 488 Tyramide and TSA™ Kit #13, with HRP-Goat Anti-Rabbit IgG and Alexa Fluor® 546 Tyramide (Life Technology, Darmstadt, Germany) were used according to the manufacturers′ instructions. For nuclear staining 1 μg/ml of DAPI (4′,6′-diamidino-2-phenylindole) (Sigma Aldrich, Taufkirchen, Germany) was used. For visualization, an Olympus AX70 microscope using cellSens software from Olympus was used.

### Ethics statement

This study has been conducted according to Declaration of Helsinki principles. IHC and immunofluorescence stainings of anonymized tissue samples as well as analysis of serum samples of cervical cancer patients were approved by the Ethics Committees of the Medical Faculty of the Saarland University at the Saarland Ärztekammer (Saarbrücken, Germany). Written informed consent was provided by all study participants.

### Succinate detection

For evaluation of succinate levels in the serum of 59 cervical cancer patients (mean age of 53.5 ± 12.9) with squamous cell carcinomas and 38 female age-matched controls (mean age of 50.4 ± 9.6) the Succinate Colorimetric Assay Kit (Sigma Aldrich) was used according to the manufacturers′ instruction.

### Tissue specimens, IHC, and IF analysis

Formalin-fixed paraffin-embedded anonymized lesions of the cervix uteri from 52 patients were taken from the local pathology archive of the Saarland University Medical Center (Homburg, Germany). The tumors were staged according to FIGO classification by expert pathologists (Y.-J. Kim or R.M. Bohle). Lesions were stained for SDHD expression and costained for CD4 and IL-17. Two-micrometer thick sections fixed on microslides were deparaffinized with xylene and hydrated using a diluted alcohol series. For immunohistochemistry or immunofluorescence, sections were incubated with citrate buffer (10 mM, pH 7.0) or with TE buffer (10 mM Tris, 1 mM EDTA, pH 9.0), respectively and cooked for 10 min or 20 min, respectively, for antigen retrieval and immersed in 3 % H_2_O_2_ in TBS (50 mM Tris/HCl, 150 mM NaCl, pH 7.6) to block endogenous peroxidase activity. To reduce nonspecific staining, each section was treated with 2.5 % normal horse serum for 30 min. Rabbit anti SDHD antibody (1:200, AB_2717499; Fisher Scientific), mouse anti-CD4 monoclonal antibody 4B12 (1:1000, Leica Biosystems; AB_563560), and rabbit anti-IL-17 polyclonal antibody (1:500, Abcam; AB_1603584) were used. For immunohistochemistry, ImmPRESS Detection Kit (Vector Laboratories, Burlingame, USA) was used. For immunofluorescence TSA^TM^ Kit #2, with HRP-Goat Anti-Mouse IgG and Alexa Fluor ^TM^ 488 Tyramide and TSA^TM^ Kit #13, with HRP-Goat Anti-Rabbit IgG and Alexa Fluor® 546 Tyramide (Life Technology, Darmstadt, Germany) were used according to the manufacturer‘s instructions. Lesions were scanned with standardized settings using Olympus BX51 microscope and VIS (Visiopharm Integrator system, Hørsholm, Denmark), Cell Sens Dimension and Microsoft Image Composite Editor Program. To evaluate the number of infiltrating Th17 cells, the numbers of CD4^+^ and IL-17^+^ cells were counted and referred to numbers/mm^2^. SDHD staining intensity was classified using the immunoreactive score (IRS; Supplementary Table S2; refs. (52)).

### Statistics

All statistical analyses were performed using the GRAPHPAD Prism 8 (GRAPHPAD Software, RRID:SCR_000306) program. To evaluate the statistical differences between the analyzed groups, a two-tailed Mann-Whitney U-test was applied for the comparison of nonparametric data between two groups and the Kruskal–Wallis test for comparison of nonparametric data of > 2 groups. Significances are indicated by asterisks (*P < 0.05; **P < 0.01; ***P < 0.001; ****P < 0.0001). Correlation between IRS of SDHD and the number of Th17 cells was done using Spearman rank correlation. Best cutoffs to discriminate patients with increased Th17 frequencies and diminished SDHD expression and with or without recurrent cervical cancers were identified by receiver operator characteristics (ROC) analysis and Youden’s index calculation.

## Acknowledgments

The authors thank all blood donors for providing their blood. The authors are grateful to Dr. Wilhelm Walch for assistance in recruitment of probes from healthy controls and Dr. Christian Herr for assistance in microscopy and slide scanning. This work was supported by a grant from the ‘Else Kröner-Fresenius Stiftung’ (2017-A64) and a grant of the ‘Wilhelm Sander-Stiftung’ (2022.012.1) to B. Walch-Rückheim.

## Author contributions

MP, MH and BWR designed the study; MP, SG, TT, MH and BWR designed the experiments;

MP, SG, LT, TT, MH and BWR developed methodology and performed experiments;

RS, NL, YJK, RMB, EM, EFS and BWR contributed to study design, patient recruitment, technical and material support and clinical data acquisition.

MP, SG, LT, NL, MH and BWR contributed to analysis and interpretation of data.

YJK contributed to conducting, analysis, and interpretation of histopathologic studies.

MP and BWR supervised all parts of the study.

BWR wrote the original draft of the manuscript. BWR funding acquisition.

MP, SG, RS, LT, NL, YJK, EFS, EM, MH and BWR involved in review and editing of the manuscript.

All authors approved the final version of the manuscript.

## Data availability

The data generated in this study are available within the article and its Supplementary data files.

