## Supplementary Figure S1-S5 for "Th17 cells target the metabolic miR-142-5p-SDHC/SDHD axis promoting invasiveness and progression of cervical cancers"

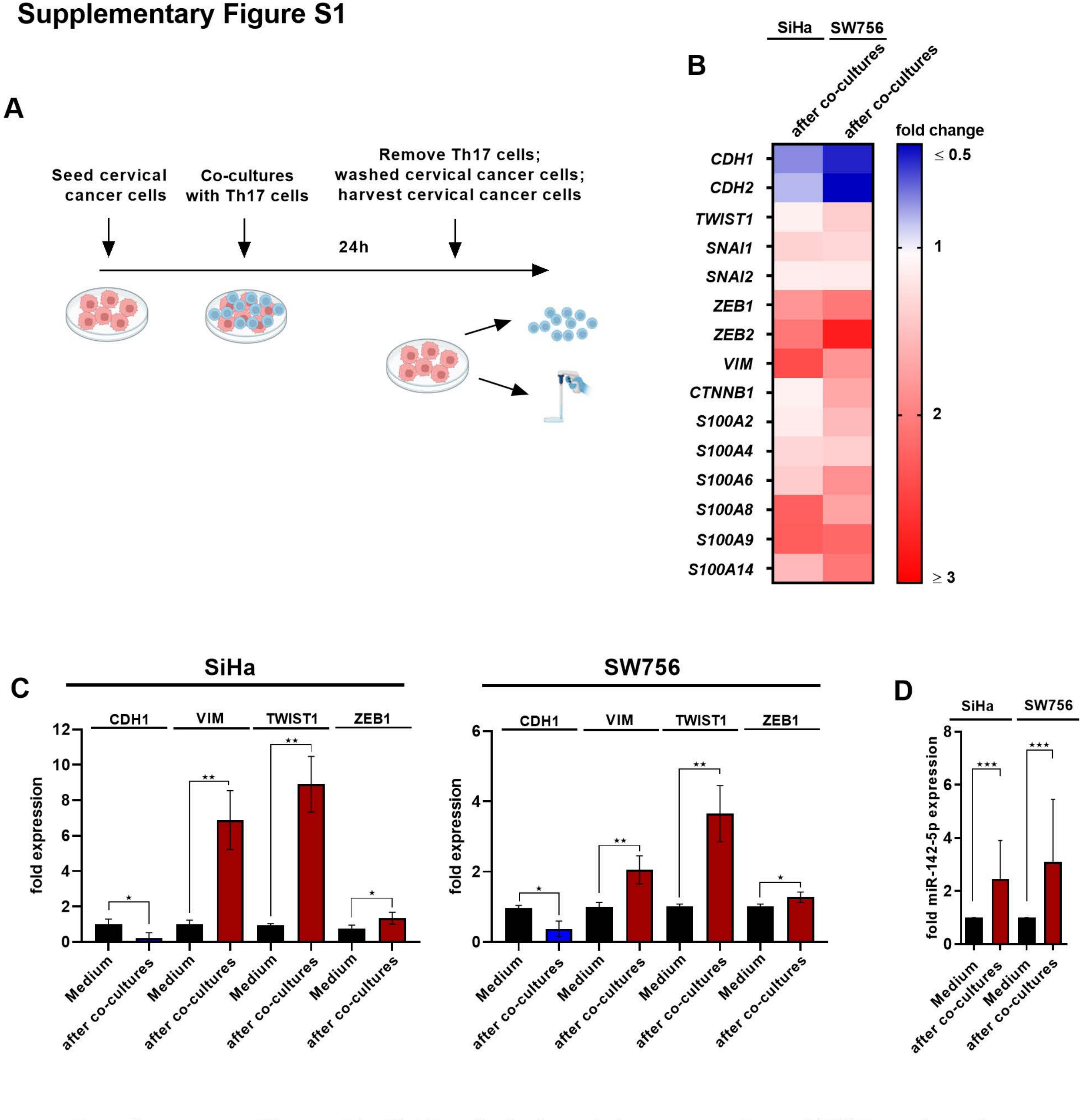


**Supplementary Figure S1: Th17 cells induced the expression of EMT markers in co-cultured cervical cancer cells.** (A) Scheme of the experimental procedures of co-cultures experiments. (B) SiHa and SW756 cells were co-cultured with Th17 cells for 24 h. Transcriptomic profile of EMT markers. Fold changes of stimulated cells to unstimulated cells were illustrated by color code (reduction in blue, induction in red). (C, D) Expression profile of EMT markers (C) or miR-142-5p expression (D) was validated by qRT-PCR. P-value according to the nonparametric Mann-Whitney U-test. Asterisks represent statistical significances: *P < 0.05; **P < 0.01; ***P < 0.001.

### Supplementary Figure S2

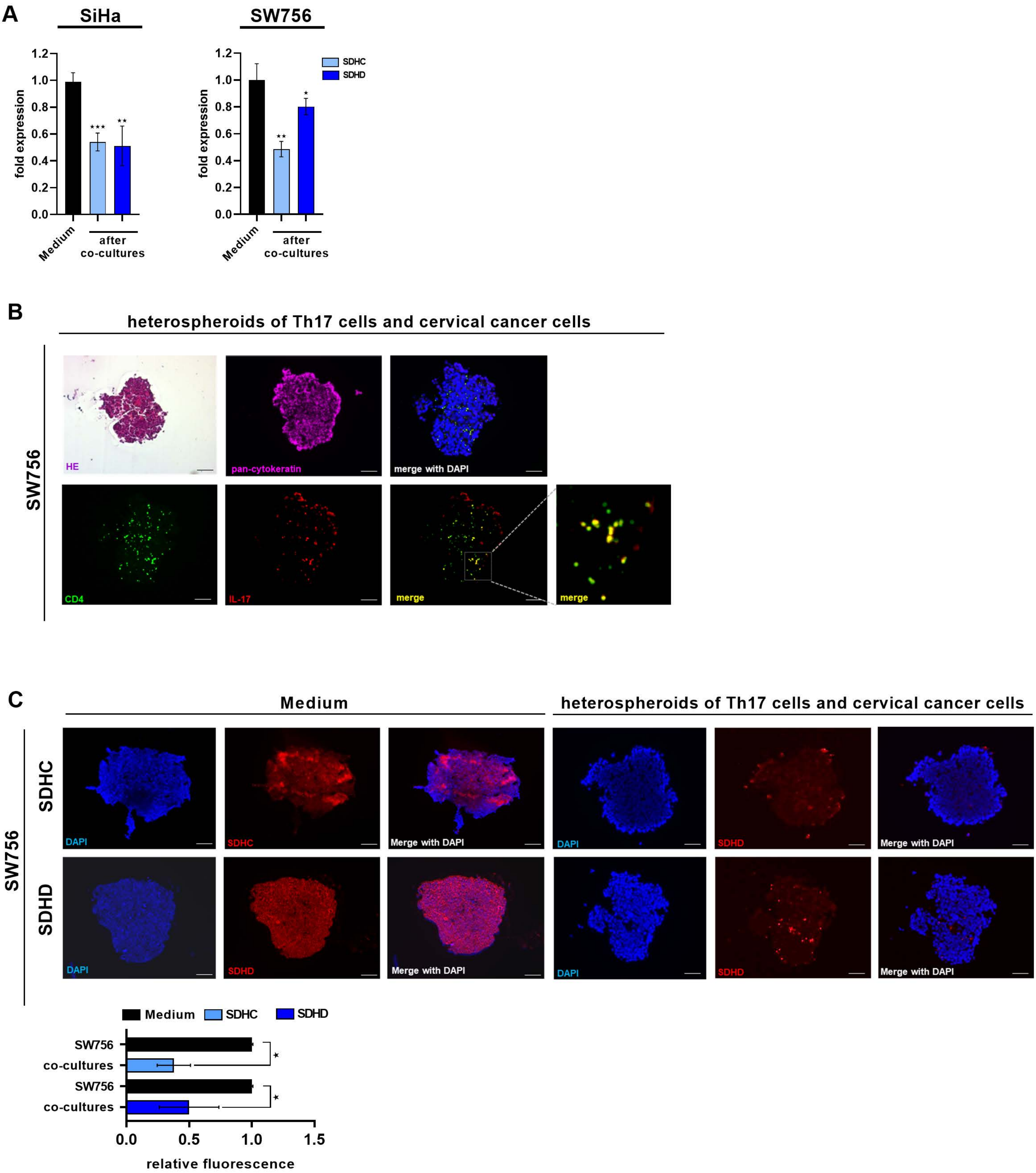


**Supplementary Figure S2: Th17 cells reduced the expression of SDHC and SDHD in cervical cancer cells.** (A) SiHa and SW756 cells were co-cultured with Th17 cells or stimulated with medium for 24 h. Expression of SDHC and SDHD was evaluated by qRT-PCR. 3D spheroids of SW756 and Th17 cells cells generated over 10 days. 5 µm sections of fixed paraffin-embedded spheroids were validated by HE stainings and analyzed for (B) CD4 and IL-17, (C) SDHC and SDHD expression by IF. Bars represent quantification of relative fluorescence/spheroid of SDHC and SDHD expression of n=6 independent spheroids, respectively. P-value according to the nonparametric Kruskal-Wallis or Mann-Whitney U-test. Asterisks represent statistical significances: *P < 0.05; **P < 0.01; ***P < 0.001.

#### Supplementary Figure S3

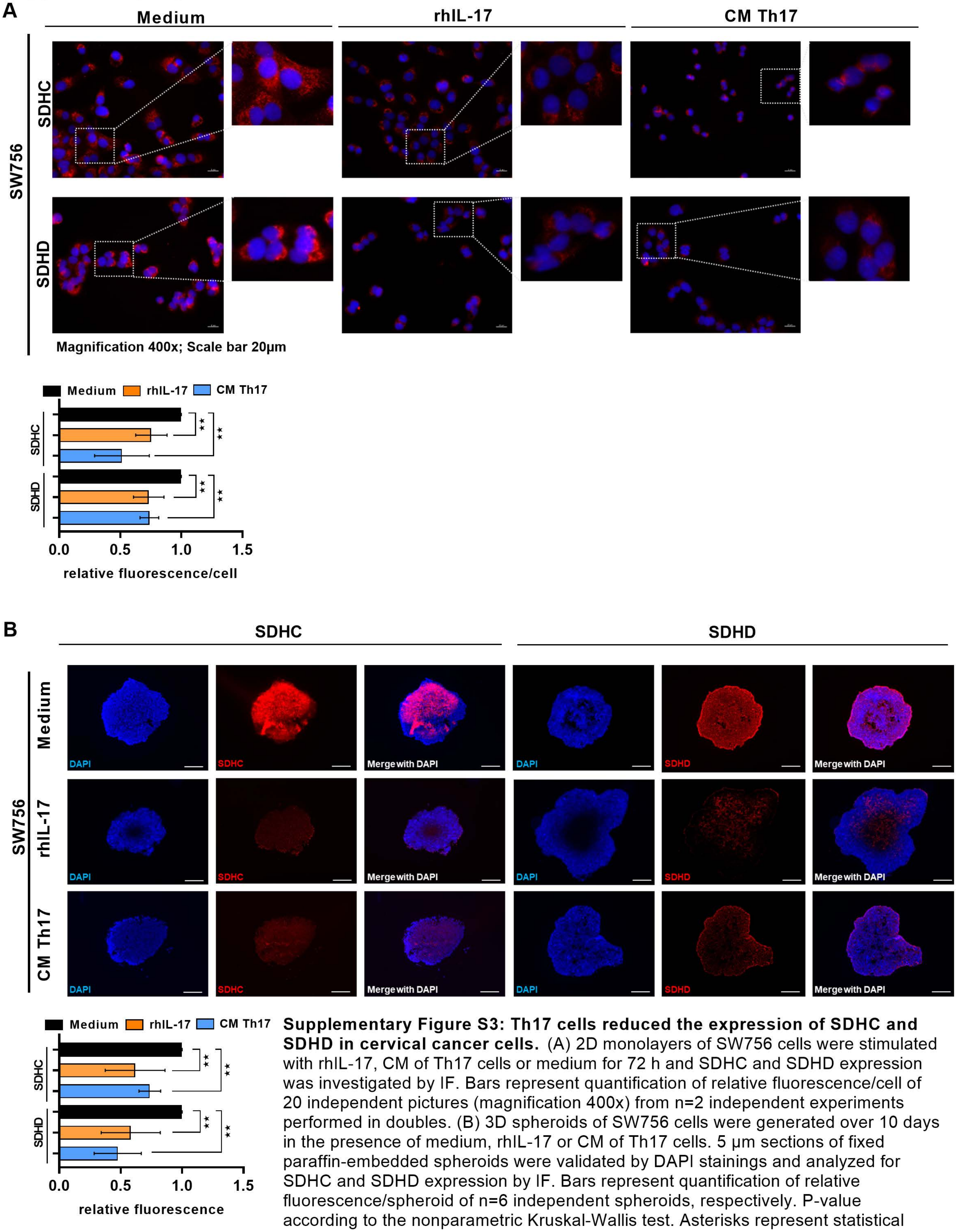


**Supplementary Figure S3: Th17 cells reduced the expression of SDHC and SDHD in cervical cancer cells.** (A) 2D monolayers of SW756 cells were stimulated with rhIL-17, CM of Th17 cells or medium for 72 h and SDHC and SDHD expression was investigated by IF. Bars represent quantification of relative fluorescence/cell of 20 independent pictures (magnification 400x) from n=2 independent experiments performed in doubles. (B) 3D spheroids of SW756 cells were generated over 10 days in the presence of medium, rhIL-17 or CM of Th17 cells. 5 µm sections of fixed paraffin-embedded spheroids were validated by DAPI stainings and analyzed for SDHC and SDHD expression by IF. Bars represent quantification of relative fluorescence/spheroid of n=6 independent spheroids, respectively. P-value according to the nonparametric Kruskal-Wallis test. Asterisks represent statistical significances: *P < 0.05; **P < 0.01; ***P < 0.001; ****P < 0.0001.

### Supplementary Figure S4

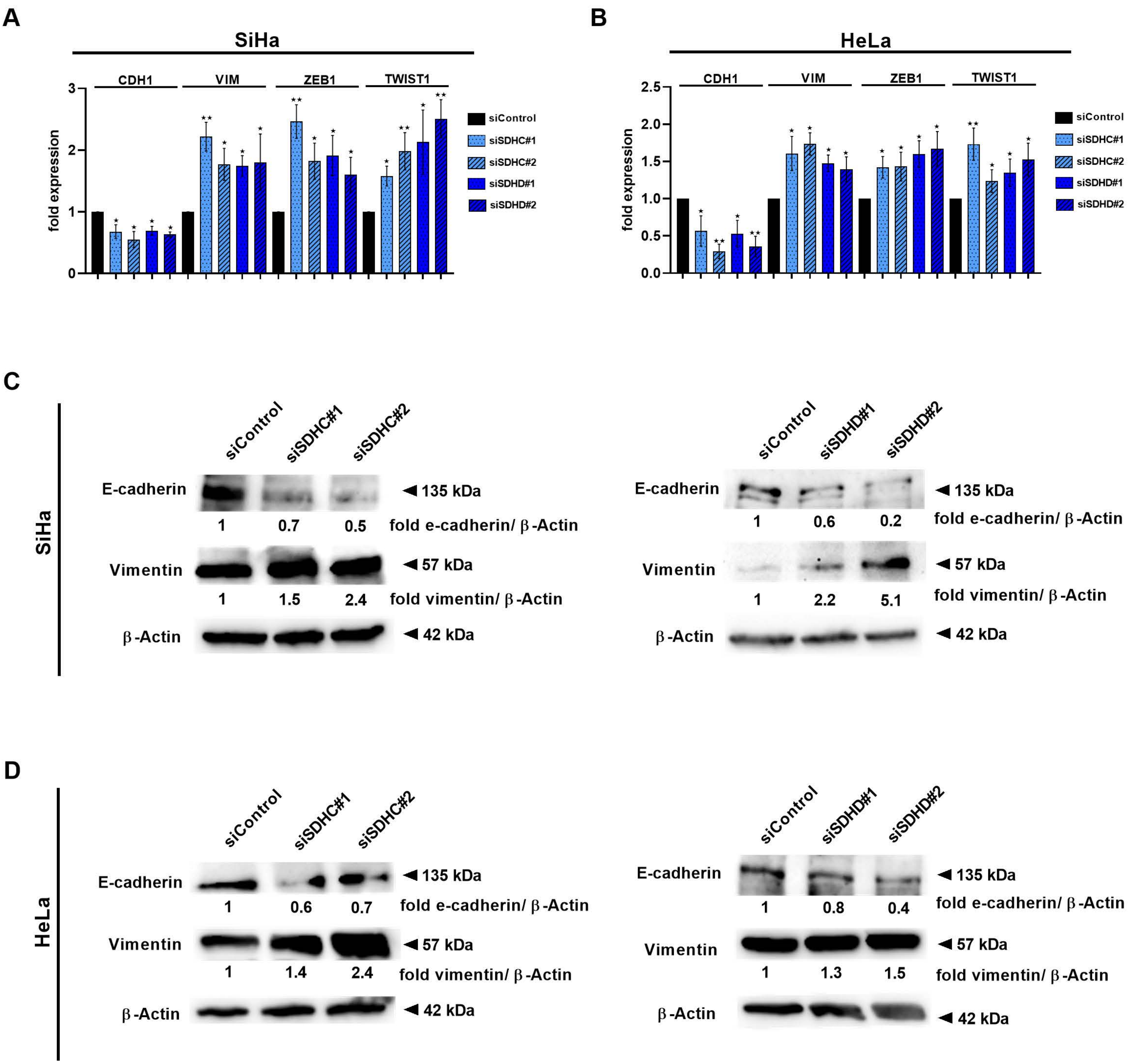


**Supplementary Figure S4: Knock down of SDHC and SDHD induced an EMT phenotype in cervical cancer cells.** SiHa and HeLa cells were transfected with two specific siRNAs for SDHC (dotted and stripped light blue bars) or SDHD (dotted and stripped dark blue bars), respectively, or mock siRNA (black bars) as a control. (A, B) After 48 h, CDH1, VIM, ZEB1 and TWIST1 expression was calculated by qRT-PCR and (C, D) e-cadherin and vimentin expression by western blot analysis. Shown are the results from n=3 independent experiments performed in triplicates in A and B and from n=2 independent experiments in C and D. Asterisks represent statistical significances: *P < 0.05; **P < 0.01.

#### Supplementary Figure S5

A

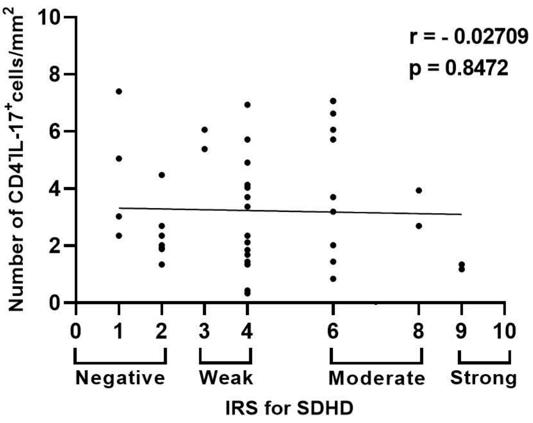


B

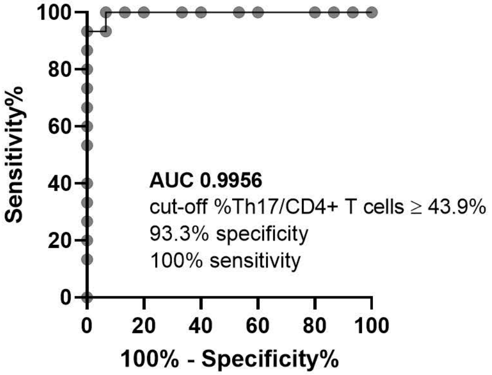


C

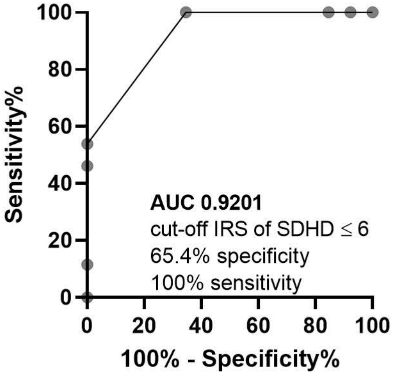


**Supplementary Figure S5: $CD4^-IL-17^+$ cells did not correlate with SDHD expression.** (A) Sections of human SCCs of 52 cervical cancer patients were stained for SDHD expression by IHC and CD4 and IL-17 by immunofluorescence. Numbers of $CD4^-$ $IL-17^+$ cells $/mm^2$ were evaluated. Correlation between $CD4^-$ $IL-17^+$ cells $/mm^2$ and IRS of SDHD. (B, C) ROC analysis of (B) percentages of Th17/$CD4^+$ T cells and (C) IRS of SDHD.
